# Independence of Centromeric and Pericentromeric Chromatin Stability on CCAN Components

**DOI:** 10.1101/2024.12.18.628896

**Authors:** Ronald J. Biggs, Kousik Sundararajan, Mingxuan Sun, Eline Hendrix, Aaron F. Straight, John F. Marko

## Abstract

The chromatin of the centromere provides the assembly site for the mitotic kinetochore that couples microtubule attachment and force production to chromosome movement in mitosis. The chromatin of the centromere is specified by nucleosomes containing the histone H3 variant CENP-A. The constitutive centromeric-associated network (CCAN) and kinetochore are assembled on CENP-A chromatin to enable chromosome separation. CENP-A chromatin is surrounded by pericentromeric heterochromatin and bound by the sequence specific binding protein CENP-B. We performed mechanical experiments on mitotic chromosomes while tracking CENP-A and CENP-B to observe the centromere’s stiffness and the role of the CCAN. We degraded CENP-C and CENP-N using auxin-inducible degrons, which we verified compromises the CCAN via observation of CENP-T loss. Chromosome stretching revealed that the CENP-A domain does not visibly stretch, even in the absence of CENP-C and/or CENP-N. Pericentromeric chromatin deforms upon force application, stretching approximately 3-fold less than the entire chromosome. CENP-C and/or CENP-N loss has no impact on pericentromere stretching. Chromosome-disconnecting nuclease treatments showed no structural effects on CENP-A. Our experiments show that the core-centromeric chromatin is more resilient and likely mechanically disconnected from the underlying pericentromeric chromatin, while the pericentric chromatin is deformable yet stiffer than the chromosome arms.

## Introduction

The centromere is a condensed region of the chromosome that plays a crucial role in chromosome segregation by providing the kinetochore assembly site (Nagpal and Fierz, 2021). The kinetochore consists of the Constitutive Centromere Associated Network (CCAN) at the inner kinetochore and the Knl1/Mis12/Ndc80 (KMN) network at the outer kinetochore (Musacchio and Desai, 2017; Hara and Fukagawa, 2018; Cairo and Lacefield, 2020; Kixmoeller, Allu, and Black, 2020). Human centromeres assemble on highly repetitive α-satellite DNA by recruiting CENtromere Proteins (CENPs) (Masumoto *et al*., 1989; Cooke, Bernat, and Earnshaw, 1990; Nagpal and Fukagawa, 2016; Musacchio and Desai, 2017; Kixmoeller, Allu, and Black, 2020). The general schematic of the centromere, specific complexes of the CCAN, and the KMN network is shown in Fig. 1 based on the structural data and models of (Musacchio and Desai, 2017; Pesenti *et al*., 2022; Yatskevich, Barford, and Muir, 2023).

**Figure 1.**
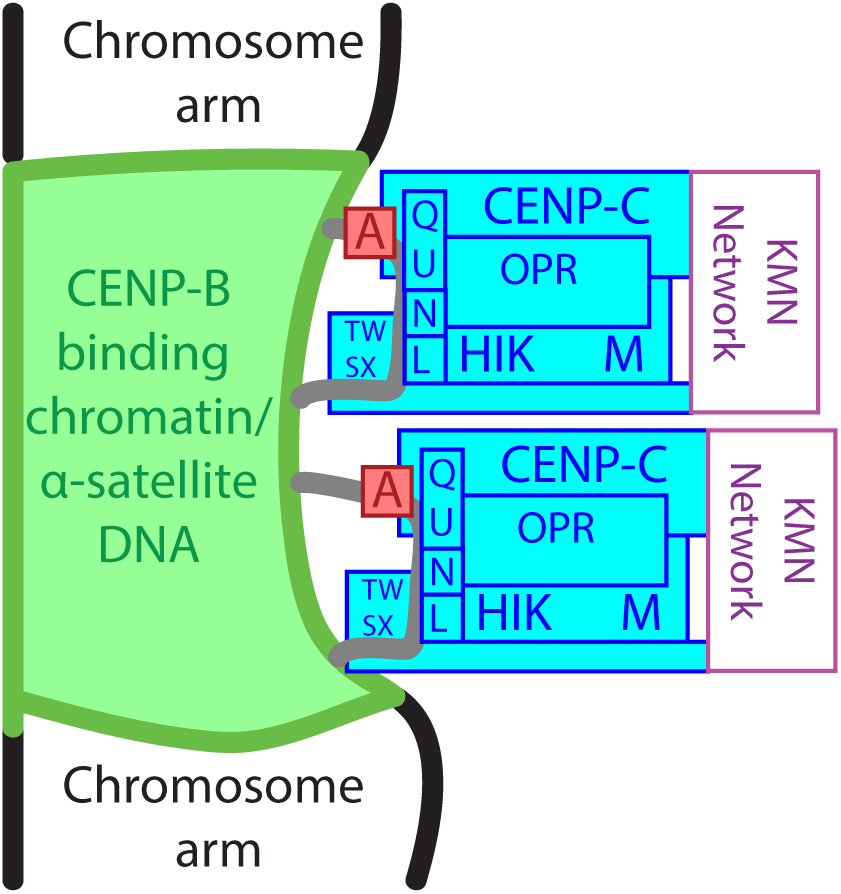
Model of the relative positioning of the centromere and associated complexes. Components of the CCAN (inner kinetochore, blue) connect the underlying pericentromeric chromatin (green) to the outer kinetochore (purple). CENP-A nucleosomes (red) are associated with CENP-C, with adjacent DNA (gray) contacting CENP-N/L and CENP-TWSX.

Nucleosomes containing the centromere specific histone H3 variant CENP-A assemble on α-satellite DNA, although not to a specific DNA sequence, where they are interspersed with the typical histone H3-containing nucleosomes (Vissel and Choo, 1987; Waye and Willard, 1989; Warburton *et al*., 1997; Blower, Sullivan, and Karpen, 2002; Sullivan and Karpen, 2004; Kixmoeller, Allu, and Black, 2020). CENP-B, a centromere protein found in the heterochromatic pericentromere, binds directly to “B-box” sequences in α-satellite DNA repeats (Cooke, Bernat, and Earnshaw, 1990). CENP-B is broadly distributed within the CENP-A-containing region of the centromere and the surrounding heterochromatin, and supports the incorporation and stability of CENP-A nucleosomes into the centromeric chromatin (Masumoto *et al*., 1989; Okada *et al*., 2007; Fachinetti *et al*., 2015; Fujita *et al*., 2015; Otake *et al*., 2020; Nagpal and Fierz, 2021).

CENP-C plays an important role in the CCAN by linking centromeric chromatin to the KMN network by directly binding CENP-A nucleosomes and proteins in the KMN network (Carroll *et al*., 2009; Carroll, Milks, and Straight, 2010; Klare *et al*., 2015). The CENP-L/N complex is also able to directly bind to CENP-A nucleosomes and can bridge adjacent nucleosomes in chromatin through nonspecific DNA binding (Carroll *et al*., 2009; Pentakota *et al*., 2017; Tian *et al*., 2018; Zhou *et al*., 2022). When assembled into the CCAN, CENP-L/N does not directly contact CENP-A but instead forms a vault or tunnel that makes extensive DNA contacts (Hara and Fukagawa, 2017; Zhang, Bellini, and Barford, 2020; Yatskevich, Barford, and Muir, 2023). CENP-C and CENP-N also help stabilize CENP-A-containing nucleosomes *in vitro* but seem to have little effect in CENP-A-containing nucleosomes in living cells (Cao *et al*., 2018). CENP-C and CENP-N directly interface with CENP-A in the CCAN and are required for CCAN stability (Klare *et al*., 2015; Hoffmann *et al*., 2016). CENP-C and CENP-N depletion has also been shown to have a myriad of negative effects cellular function, including cellular lethality (Howman *et al*., 2000; Kwon *et al*., 2007; McClelland *et al*., 2007; McKinley *et al*., 2015).

During mitosis, large microtubule/motor-based forces are applied to the KMN network and transduced through the CCAN to the centromeric chromatin (Kajtez *et al*., 2016; Ye, Cane, and Maresca, 2016; Maiato *et al*., 2017; Anjur-Dietrich, Kelleher, and Needleman, 2021). Each human centromere experiences spindle forces on the order of hundreds of pN, applied by roughly 9 microtubules (Kiewisz *et al*., 2022). During merotelic attachments (a microtubule attachment defect where one kinetochore is attached to both spindle poles), the centromere can be pulled apart (observed via CREST staining) suggesting that the centromere can be stretched by spindle forces applied directly across it (Cimini *et al*., 2001). Additionally, CENP-A-stained centromeres can periodically be seen deforming in normal attachments, but only when missing CENP-C (Suzuki *et al*., 2014).

We hypothesized that the centromeric chromatin, stabilized by interactions between the CCAN, CENP-A, and CENP-B must be quite strong to survive spindle forces during mitosis. While stabilization experiments typically focus on the premature separation of sister centromeres when the CCAN is perturbed, we studied the mechanical properties of the centromere by physically stretching chromosomes along their long/longitudinal axis. We tracked the centromere and pericentromere by tracking CENP-A and CENP-B. We rapidly degraded CENP-C, CENP-N, or both using auxin-induced-degron (AID) tags to determine the role they have in promoting centromeric chromatin stiffness (Cao *et al*., 2018). CENP-A did not change in size upon stretching the whole chromosome while the CENP-B region stretched approximately one-third the amount of the whole chromosome. These behaviors did not change upon auxin-induced degradation of CENP-C or CENP-N, either alone or coincidentally. The whole CCAN was compromised by a reduction in CENP-T fluorescence following CENP-C and CENP-N degradation, either alone or coincidentally. Nuclease treatment also had no effect on the size of the CENP-A in chromosome bundles.

## Results

### AID-CENP-C, CENP-N, and CENP-C/N are degraded following auxin treatment

To determine the role of CENP-C and CENP-N in force resistance at centromeres we used YFP-AID-CENP-C and sfGFP-AID-CENP-N cell lines to both monitor the localization and amount of the proteins, before and after auxin induced degradation (Cao *et al*., 2018). Both cell lines also contained a mRuby2-CENP-A fusion protein providing a readout for the CENP-A localization and size. We also utilized a cell line containing both sfGFP-AID-CENP-C and Ruby- AID-CENP-N proteins (CENP-C/N) for their simultaneous degradation. CENP-C, CENP-N, and CENP-A localization in untreated cells showed punctate, coincident fluorescence signals in their fluorescent channels along the DNA (Hoechst staining), consistent with their centromere localization (Fig. 2A). The method of signal quantification can be seen in Figure 2B. Auxin addition reduced CENP-C and CENP-N fluorescence to the background levels of the cell (Fig. 2C) while CENP-A fluorescence persisted as punctate dots at mitotic centromeres at the same levels as the untreated centromeres (Fig. 2D). Depletion of both CENP-C and CENP-N reduced fluorescence of both proteins to background levels in the CENP-C/N line (Fig. 2C, D). Supplementary Figure 1 details the fluorescence for the two fluorescent channels as single data points with and without auxin as well as showing the two CENP-N lines’ fluorescence.

**Figure 2.**
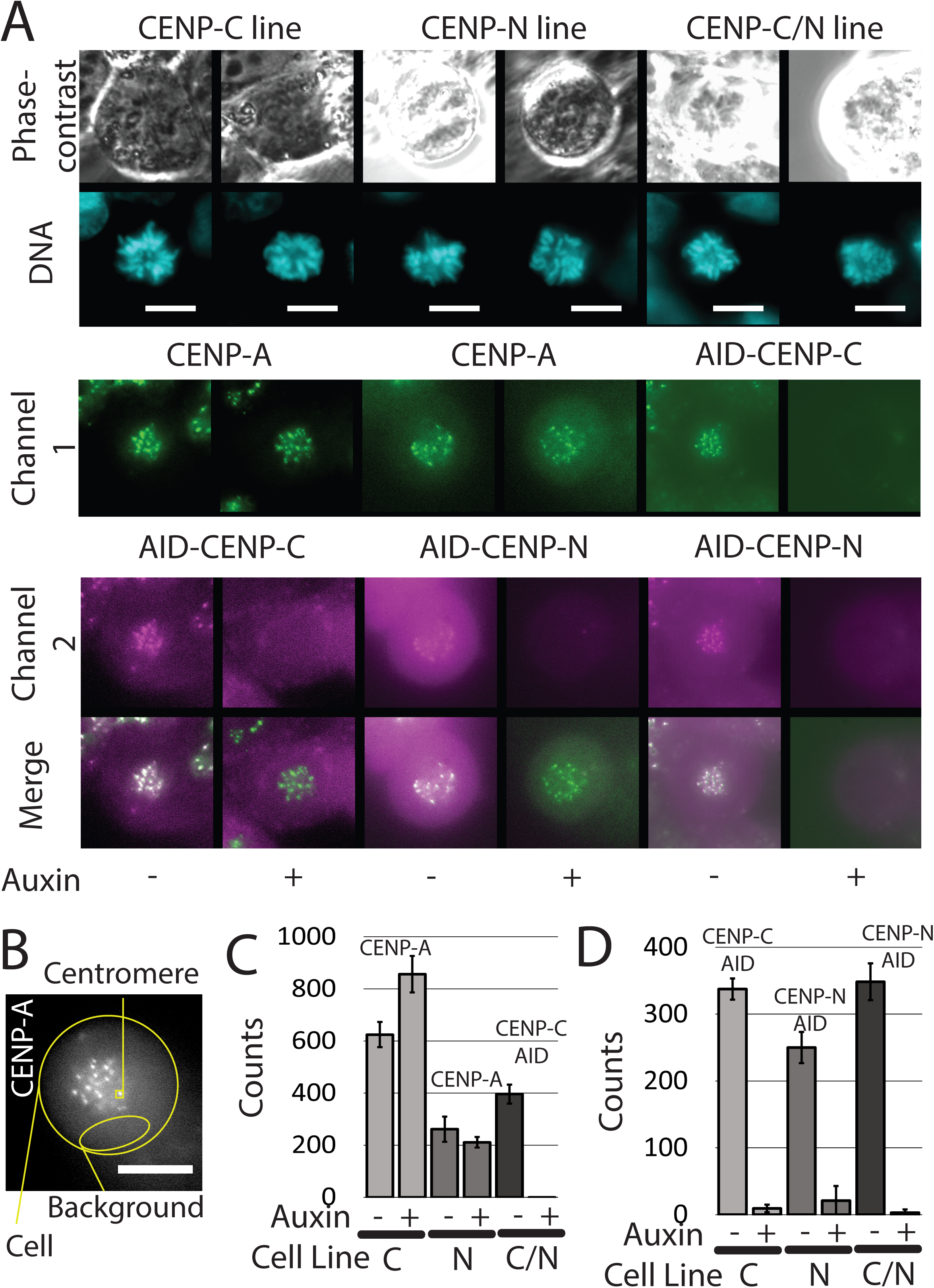
Fluorescence imaging and degradation of the used AID-CENP cell lines. (A) Example images of mitotic cells with centromere fluorescence. Top row shows phase- contrast imaging of target mitotic cells. Second row shows corresponding DNA signal via Hoechst staining in cyan (used here to show relative positions of centromere and DNA; Hoescht staining was not used in experiments that measured mechanical properties of chromosomes or the centromere). Third and fourth rows show endogenous, CENP protein in green (CENP-A or CENP-C) and magenta (CENP-C or CENP-N); labels are shown above each pair of images. Bottom row shows merged centromere images. The first two columns show the AID-CENP-C- containing line, the middle two columns show the AID-CENP-N-containing lines, and the last two columns show the AID-CENP-C and AID-CENP-N-containing line. Odd columns show untreated cells and even columns show auxin-treated cells. Auxin treatment removed the AID- tagged fluorophores’ signal but did not affect the fluorescence of non-AID-tagged CENP-A. Bar indicates 10 µm. (B) Fluorescence quantification method and example image. Representative cell showing endogenous CENP-A fluorescence with areas of interest for centromere and background intensities. In short, centromere fluorescent values are measured as the fluorescence average of the boxed centromere minus the fluorescence average of the background (inside the cell) at 500 ms exposure. Bar indicates 10 µm. (C) Quantification of fluorescence in channel 1. Numbers shown are averages of the centromere signal minus the cell background signal at 500ms exposure. In the CENP-C line, the CENP-A fluorescence was 624+/-48 counts (N=43) in untreated cells and 856+/-70 counts above background (N=20) (statistically significantly higher) in auxin-treated cells, demonstrating auxin does not have a diminishing effect on the CENP-A signal. In the CENP-N (409) line, the CENP-A fluorescence was 261+/-48 counts above background (N=13) and 211+/-21 counts (N=12) (not statistically different) in auxin-treated cells. In the CENP-C/N line, CENP-C fluorescence was 396+/-37 counts above background (N=26) and -7+/-2 counts above background (N=21) (statistically significantly lower) in 4-hour auxin-treated cells. Single data points corresponding to these averages are shown in Supplementary Figure 1A. (D) Quantification of auxin degradation (CENP-C, CENP-N (409), and CENP-N respectively). In the CENP-C line, the CENP-C fluorescence was 338+/-16 counts above background (N=42) in untreated cells and 9+/-6 counts above background (N=20) (statistically significantly lower) in auxin-treated cells. In the CENP-N line, the CENP-N fluorescence was 208+/-33 counts above background (N=26) in untreated cells and 21+/-22 counts above background (N=24) (statistically significantly lower) in auxin-treated cells. In the CENP-C/N line, the CENP-N fluorescence was 348+/-28 counts above background (N=26) in untreated cells and 3+/-5 counts above background (N=21) (statistically significantly lower) in 4-hour auxin-treated cells. Single data points are shown in Supplementary Figure 1B.

### Overall chromosome mechanics are unaltered by CENP-C/N degradation

To determine the effects of CENP-C and CENP-N degradation on chromosome mechanics, we measured the doubling force of the isolated chromosomes by stretching the chromosomes in their linear, elastic-response regime (under a 4-fold length increase, Fig. 3 A-C; Supplementary Fig. 2A, first three panels). We found that removal of CENP-C, CENP-N, or both caused no statistically significant change to the doubling force or elastic modulus of the whole chromosome for any of the cell lines studied (Supplementary Fig. 2B).

**Figure 3.**
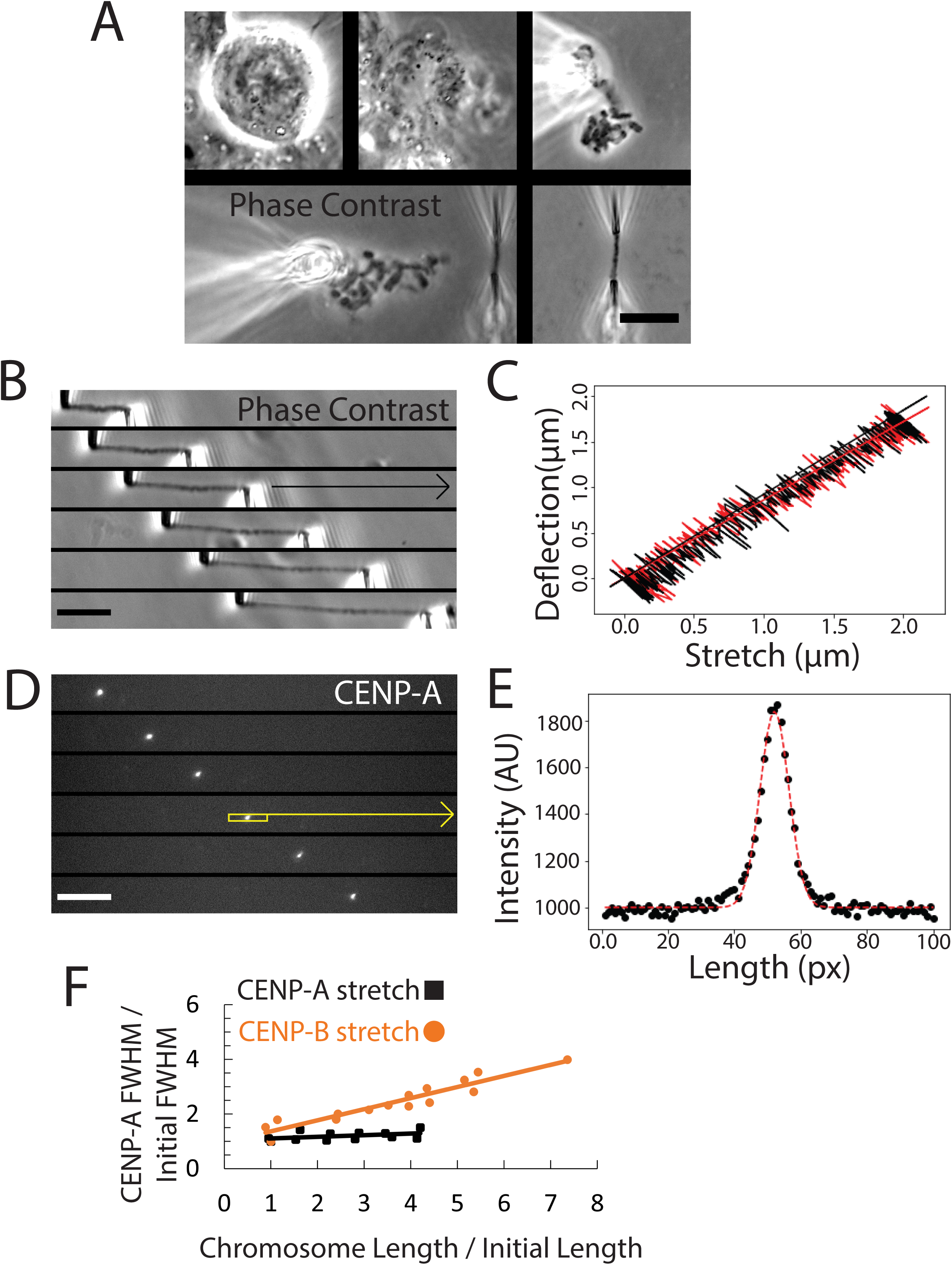
Methods for chromosome isolation, mechanics, and quantification. (A) Example of chromosome isolation by bundle removal method. Upper left- Target mitotic cell. Upper middle- Target cell after lysis by detergent. Upper right- Chromosome bundle after aspiration into stabilizing pipette. Lower left- Target chromosome aspirated into chromosome- aspirating pipettes. Lower right- Isolated target chromosome. Scale bar 10 µm long, lower right. (A) Example experiment stretching a whole chromosome in phase-contrast imaging. (C) Chromosomes were stretched by moving the pull pipette (bottom) away from the deflecting pipette (top). An example of a live stretch-deflection trace is shown in (C), while the corresponding fluorescent images are shown in (D). The chromosome vs centromere stretch is plotted in (F), where the chromosome stretch is manually measured as the distance between the middle of the pipettes. Bar represents 10 µm. (B) Example trace of a stretch-deflection experiment for a whole chromosome. A computer tracked the two pipettes’ positions simultaneously through the experiment and directed a micromanipulator to move the pull pipette away from the deflecting pipette at a specified speed and distance (typically 1.0 µm/sec and 20 µm respectively in these experiments). The linear regression slope of chromosome stretching vs pipette deflection was multiplied by the spring constant of the deflection pipette to obtain the spring constant of the whole chromosome. Supplementary Figure 2A shows an additional force-extension plot, including past its linear- stretching regime and Supplementary Figure 2B shows the individual data points of the mechanics of all chromosomes. (C) CENP-A fluorescence during chromosome stretching. CENP-A images were taken while stretching the chromosome (B). The yellow box was placed around the centromere and averaged over each column of the box’s length and plotted in (E) to obtain its length. Bar indicates 10 µm. (D) Full width at half maximum (FWHM)/fluorescent trace of CENP-A. Black dots represent the raw data of the box plot seen in (D). Red dashed line shows a gaussian fit, used to calculate FWHM, giving the centromere length in pixels. The FWHM was converted from pixels to microns when reporting CENP-A lengths and corrected for its point-spread function. This was repeated for each image of the centromere and was plotted against chromosome stretch in (F). (E) Plot of centromere stretch vs whole-chromosome stretch. Chromosome length (B) was plotted against centromere FWHM (D, E), both normalized to initial unstretched chromosome length and centromere FWHM. A linear fit gave a slope of 0.057+/-0.046 for the CENP-A stretch (black line and black squares). Another chromosome stretched with CENP-B staining is also shown in the graph plotted against chromosome stretch. A linear fit yielded slope of 0.407+/-0.038 (orange line and orange circles). Slopes were determined for each centromere and CENP-B stretching experiment (Fig. 4B, D and 5B; one data point for each average of a series of individual chromosome stretches is shown in Supplementary Fig. 3).

### The CENP-A region does not observably stretch during whole-chromosome stretching in the presence or absence of CENP-C, CENP-N or both

We then stretched the whole chromosome while tracking the endogenous CENP-A fluorescence (Fig. 3D-F, 4A) and observed no extension of the CENP-A region (Fig. 4B, left panel), even for large chromosome deformations, including past its linear-response regime (Supplementary Fig. 2A, last panel). Quantification of centromere and chromosome length showed that in untreated cells, the centromere, via CENP-A tracking, did not stretch or deform, while the whole chromosome stretched several-fold its initial length (Fig. 3B-F, 4A (examples), B (left panel) (average)). The ratio of centromere: chromosome stretch in the whole-chromosome- stretching experiments showed an average slope near zero consistent with no centromere deformation within the error range of our measurements (Fig. 4B, left panel).

**Figure 4.**
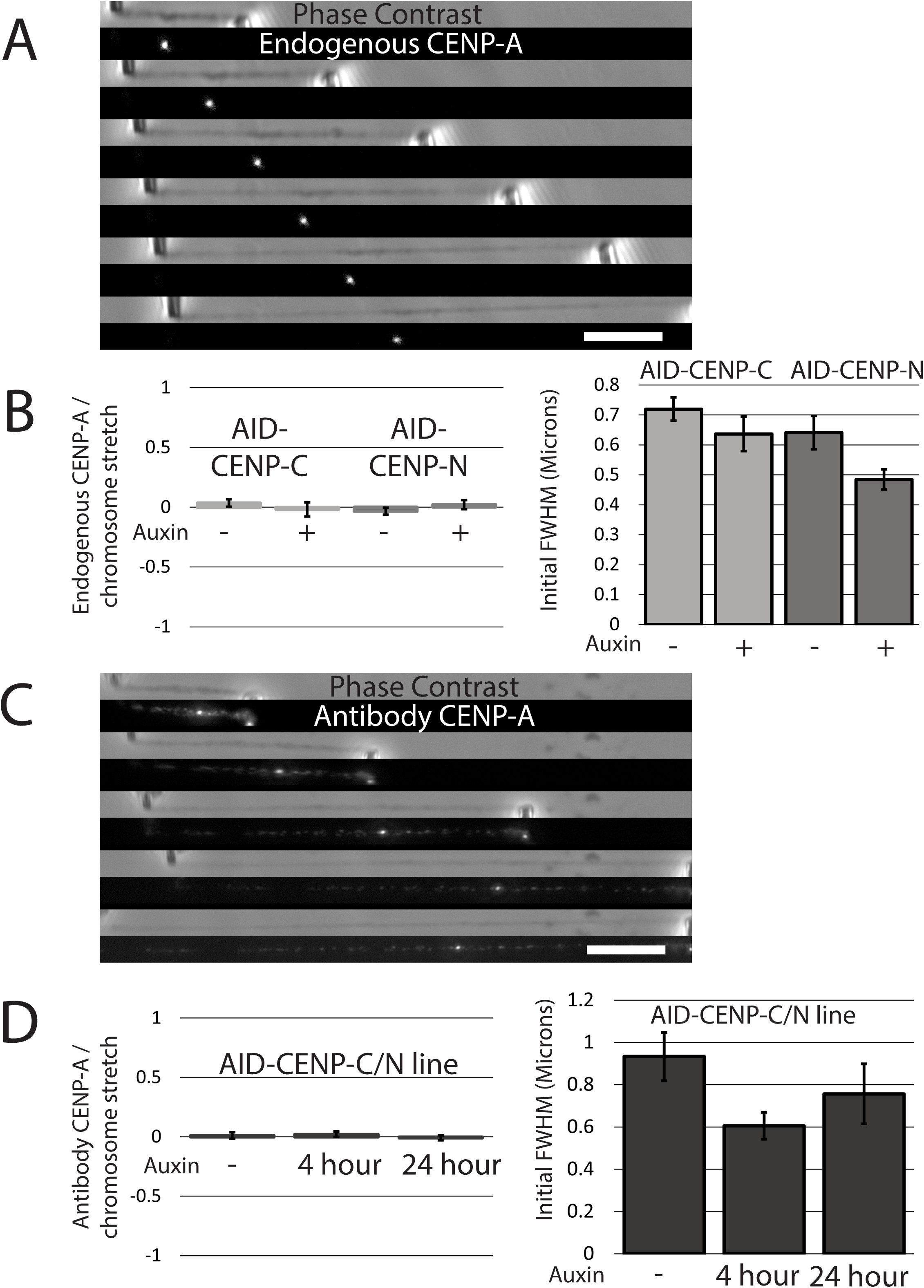
CENP-A does not stretch but may become smaller when CENP-C or N is degraded. (A) Example chromosome stretch showing centromere behavior (endogenous CENP-A). The whole chromosome in phase-contrast imaging is shown above each image of the endogenous centromere in its corresponding fluorescent channel. The example image shows a CENP-A stretch ratio of 0.058 (centromere stretch: chromosome stretch) with an initial length of 0.59μm after point-spread correction. Bar indicates 10 µm. (B) Quantification of endogenous CENP-A mechanics dependence on CENP degradation. (C) Left panel shows the average stretch of the centromere to the whole chromosome. In CENP-C cells, the endogenous CENP-A: chromosome stretch ratio was 0.035+/-0.032 (N=28) in untreated cells, and -0.018+/-0.058 (N=14) (statistically insignificantly different) in auxin-treated cells. In CENP-N cells, the endogenous CENP-A: chromosome stretch ratio was -0.033+/-0.058 (N=14) in untreated cells, and -0.022+/-0.039 (N=15) (statistically insignificantly different) in auxin-treated cells. The individual data corresponding to these averages are shown in the first four columns of Supplementary Figure 2A. The right panel shows the average of the initial length of the centromere (endogenous fluorescent CENP-A signal) (the centromere’s length when the chromosome is at its initial length and under no tension). Average lengths are corrected for the microscope fluorescence point-spread function (Supplementary Fig. 7A, B). The initial length of the endogenous CENP-A in the CENP-C line was 0.71+/-0.04 µm (N=41) in untreated cells, and 0.63+/-0.07 µm (N=14) (smaller, but not statistically different) in auxin-treated cells. The initial length of the endogenous CENP-A in the CENP-N line was 0.63+/-0.06 µm (N=18) in untreated cells, and 0.48+/-0.04 µm (N=14) (marginally statistically significantly lower (P=0.054)) in auxin-treated cells. The single data points can be seen in the first four columns of Supplementary Figure 2B. (C) Example stretching of a chromosome with the centromere (antibody-labeled CENP-A). The whole chromosome in phase-contrast imaging is shown above each image of the antibody-stained centromere in its corresponding fluorescent channel. The example image shows a CENP-A stretch ratio of -0.045 (centromere stretch: chromosome stretch) with an initial length of 0.36μm after point-spread correction. While there is non-specific binding, as seen by the frequency of fluorescent signal along the chromosome, an increase in fluorescence can be seen in a punctate area, corresponding to the centromere, used in the quantification shown in Figure 4D. Bar indicates 10 µm. (D) Quantification of antibody CENP-A mechanics dependence on CENP degradation. The left panel shows the average stretch ratio of the centromere to the whole chromosome. In CENP-C/N cells, the antibody-stained CENP-A: centromere ratio was 0.011+/-0.028 (N=16) in untreated cells, 0.022+/-0.024 (N=17) (statistically insignificantly different from all treatments) in 4-hour auxin-treated cells, and -0.007+/-0.020 (N=7) (statistically insignificantly different from all treatments) in 24-hour auxin-treated cells. Single data points are shown in the last three columns of Supplementary Figure 2A. The right panel shows the average of the initial length of the centromere (CENP-A signal after fluorescent antibody spraying) (the centromere’s length when the chromosome is at its initial length and under no tension). The reported lengths are corrected for the microscope fluorescence point-spread function (Supplementary Fig. 2). The initial length of the antibody-stained CENP-A in the CENP-C/N line was 0.91+/-0.13 µm (N=16) in untreated cells, 0.58+/-0.08 µm (N=17) (statistically significantly smaller compared to untreated) in 4-hour auxin-treated cells, and 0.62+/-0.18 µm (N=8) (statistically insignificantly different from other treatments) in 24-hour auxin-treated cells. The single data points can be seen in the last three columns of Supplementary Figure 2B.

To test the contribution of CENP-C and CENP-N to centromere stretch resistance, we measured the initial length of the CENP-A signal (*i.e.* under no stress) and the centromere: chromosome length ratio in whole-chromosome-stretching experiments after CENP-C and CENP-N degradation. After degradation of CENP-C or CENP-N via auxin treatment, we found a reduction in the initial/zero-stress length of the CENP-A region of the unstretched chromosome (Fig. 4B, right panel) (statistically insignificantly different in the CENP-C line, but a P value of 0.055 in the CENP-N line). Upon stretching the chromosomes after CENP-C or CENP-N degradation, we observed no change in the centromere to chromosome stretch ratio (as measured

by the methods detailed in Figure 3D-F) (Fig. 4B left-hand panel). The single data point measurements are shown in Supplementary Figure 3A, B.

To determine whether the simultaneous loss of both CENP-C and CENP-N impacts the mechanical response of centromeric chromatin, we simultaneously degraded both proteins using the double-degron cell line (CENP-C/N), which lacks endogenously, fluorescent-tagged CENP-A. We sprayed a fluorescent CENP-A antibody onto the isolated chromosomes, which provided a strong, localized fluorescence signal at the centromere like the endogenous CENP-A label in the single-degron cell lines (Fig. 4C). We also noticed regions of weaker fluorescence along the chromosome, which we attributed to off-target binding (Fig. 4C), but only used the high-intensity signal in measuring the length of CENP-A/core-centromeric chromatin. In chromosomes from untreated cells, the high-intensity, antibody-labeled CENP-A signal was coincident with the CENP-C and CENP-N fluorescence signals and did not lengthen upon chromosome stretching (Fig. 4D, left), consistent with the results for the single-degron cell lines (Fig. 4B).

After simultaneous degradation of both CENP-C and N via auxin treatment for 4 hours, we found it caused no change in the CENP-A: chromosome stretch ratio (Fig. 4D, left), but the initial CENP-A length was shorter (Fig. 4D, right). To test if long-term, simultaneous removal of CENP-C and CENP-N would perturb the CENP-A signal, we treated the double degron cell line with auxin for 24 hours, then analyzed the initial length and deformation of CENP-A in the stretched chromosome. We observed that there was no change in the initial length of CENP-A (the initial length was statistically insignificantly different from the untreated CENP-A and the 4-hour auxin treated CENP-A initial length due to the spread in the data) (Fig. 4D left panel, rightmost column), nor its deformation when stretching the whole chromosome (Fig. 4D, right panel, rightmost column). The single data point measurements are shown in Supplementary Figure 3A, B.

### The CENP-B region deforms while stretching the whole chromosome, but CENP-B deformation ratio does not change with CENP-C/N degradation

The result that the CENP-A/core-centromere region initial length was affected by CENP degradation, but never consistently deformed upon chromosome stretching prompted us to analyze the stretch response of the broader centromere and pericentromere. To that end, we labeled and tracked CENP-B with fluorescent antibodies (non-fluorescent primary and a fluorescent secondary antibody). We observed that the CENP-B signal localized to a small region of the chromosome around where the CENP proteins were located (Fig. 5A). When we stretched the chromosome, we observed that the CENP-B region stretched in a roughly linear fashion along with the whole chromosome in the untreated cell lines (Fig. 3F, 5A). The example slope in Figure 3F is about 0.4, indicating that chromosome’s CENP-B region was approximately 2.5-fold (1/0.4) stiffer than the whole chromosome.

**Figure 5.**
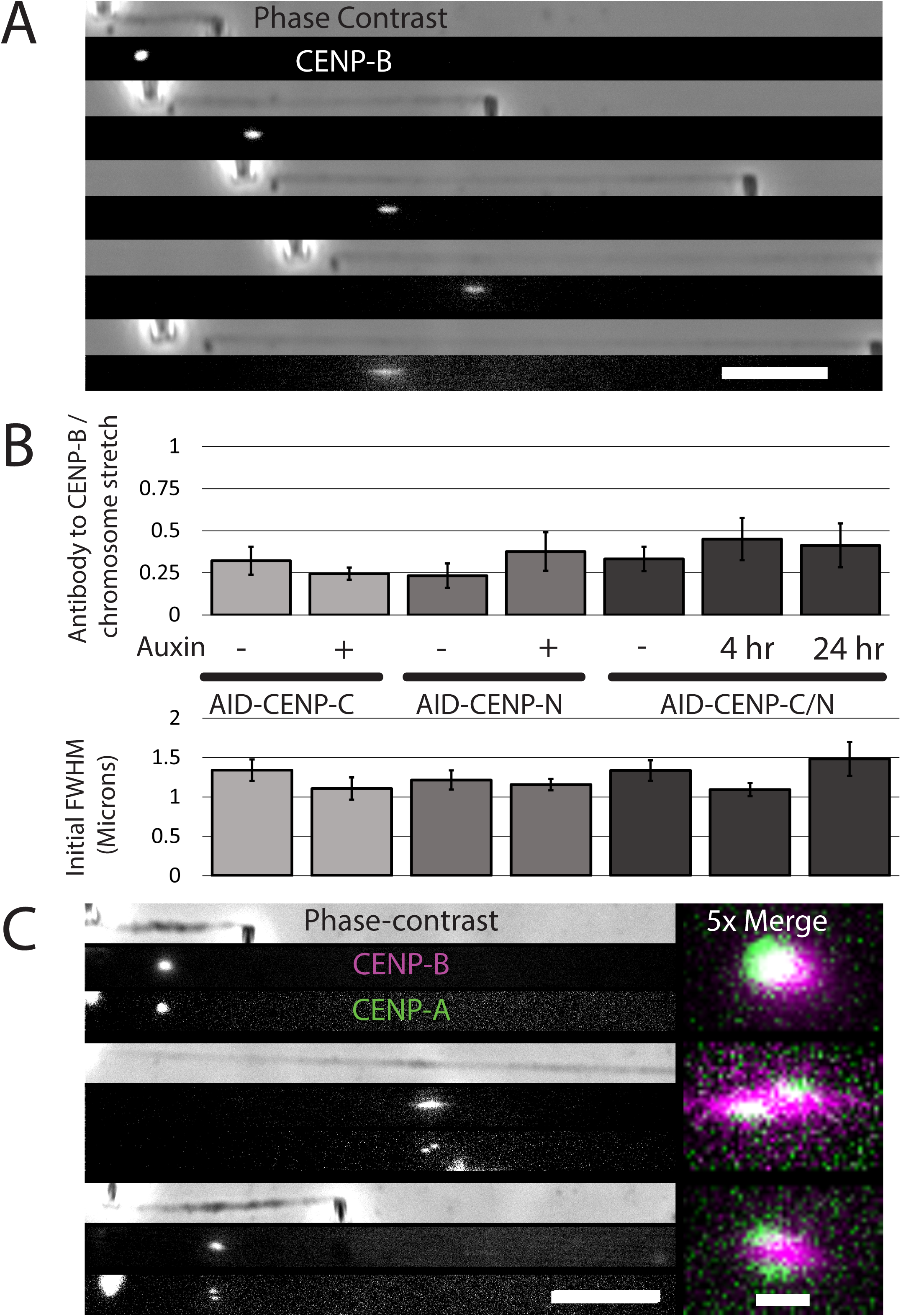
CENP-B stretches less than the chromosome, but is not affected by CENP degradation. (A) Example stretching of a chromosome with the pericentromere, antibody-labeled CENP-B. The whole chromosome in phase-contrast imaging is shown above each image of the CENP-B antibody-stained pericentromere in its corresponding fluorescent channel. The example image shows CENP-B stretching via a smear of fluorescence. The example image shows a CENP-B stretch ratio of 0.262 (centromere stretch: chromosome stretch) with an initial length of 1.20μm after point-spread correction. Bar indicates 10 µm. (B) Quantification of antibody-labeled CENP-B mechanics dependence on CENP degradation. The top panel shows the average stretch of the pericentromere to the whole chromosome. In CENP-C cells, the antibody-stained CENP-B: chromosome stretch ratio was 0.322+/-0.083 (N=8) in untreated cells, and 0.244+/-0.036 (N=10) (statistically insignificantly different) in auxin- treated cells. In CENP-N cells, the antibody-stained CENP-B: chromosome stretch ratio was 0.233+/-0.072 (N=9) in untreated cells, and 0.376+/-0.115 (N=9) (statistically insignificantly different) in auxin-treated cells. In CENP-C/N cells, the antibody-stained CENP-B: chromosome stretch ratio was 0.332+/-0.083 (N=12) in untreated cells, 0.450+/-0.072 (N=8) (statistically insignificantly different from all treatments) in 4-hour auxin-treated cells, and 0.413+/-0.131 (N=10) (statistically insignificantly different from all treatments) in 24-hour auxin-treated cells. Single data points are shown in Supplementary Figure 2C. The bottom panel shows the average initial length of the pericentromere (length of CENP-B signal after fluorescent antibody spraying when chromosome is at its initial length and under no tension). Reported lengths are corrected for the microscope fluorescence point-spread function (Supplementary Fig. 7). Initial length of the antibody-stained CENP-B in the CENP-C line was 1.34+/-0.14 µm (N=9) in untreated cells, and 1.11+/-0.14 µm (N=11) (statistically insignificantly different) in auxin-treated cells. The initial (C) length of the antibody-stained CENP-B in the CENP-N line was 1.22+/-0.12 µm (N=10) in untreated cells, and 1.16+/-0.07 µm (N=8) (statistically insignificantly different) in auxin-treated cells. The initial length of the antibody-stained CENP-B in the CENP-C/N line was 1.34+/-0.13 µm (N=12) in untreated cells, 1.09+/-0.08 µm (N=7) (statistically insignificantly different from all treatments) in 4-hour auxin-treated cells, and 1.48+/-0.21 µm (N=10) (statistically insignificantly different from all treatments) in 24-hour auxin-treated cells. The single data points are shown in Supplementary Figure 2D. (D) Example image of CENP-A localization with CENP-B signal. After spraying with CENP- B antibody in the CENP-C line, we observed colocalization of CENP-A with CENP-B. The CENP-A signal was periodically seen separating into distinct entities as each of the chromatid centromeres moved away from each other. These individual centromeres were still seen inside the CENP-B signal. This image also illustrates that even when the CENP-B signal was observed to stretch, the CENP-A signal was not seen deforming as the entire chromosome was stretched. More examples are shown in Supplementary Figure 5. Main scale bar is 10 µm, middle lower right. 5x Merge (right) is 5 times the size of the other images with CENP-A in green and CENP- B in magenta; scale bar for these images is 1 µm (lower right).

To test if loss of CENP-C or CENP-N impacted CENP-B’s domain stiffness we depleted CENP-C or CENP-N individually for 4 hours and simultaneously for 4 and 24 hours via auxin treatment. None of these treatments statistically significantly changed either deformability of CENP-B from untreated conditions (Fig. 5B, top panel) or the initial lengths (Fig. 5B, bottom panel). The individual data points can be seen in Supplementary Figure 3C, D. We observed that the CENP-B region stretched in two distinct ways: first, we observed it to separate into multiple “islands” when stretched (Supplementary Fig. 4A, B). Alternately, the CENP-B region would spread out evenly when stretched (Supplementary Fig. 4C). Both examples of stretching obeyed the proportionally linear behavior, roughly 3-fold stiffer on average than the stiffness of the whole chromosome across all cell lines (Supplementary Fig. 4).

### Relative displacements and deformations of CENP-A and CENP-B regions

We simultaneously imaged both CENP-A and CENP-B to find their relative positions and mechanical behaviors (Fig. 5C, Supplementary Fig. 5; note the latter shows several experiments to demonstrate the range of distributions). We utilized the YFP-AID-CENP-C with the mRuby2- CENP-A line due to its sufficiently high fluorescence intensity and stable CENP-A fluorescence, imaged with fluorescent antibodies against CENP-B. In the experiment shown in Fig. 5C, stretching the chromosome caused the two CENP-A signals from the two chromatids in the chromosome to separate, remaining lateral to the CENP-B signal, which can also be seen in Supplementary Figure 5. For untreated cells, the CENP-B signal appeared close and connected to the CENP-A region signal(s) as the chromosome stretched (Supplementary Fig. 5A-B). We observed that there may be more separation (loss of coincident signals) of CENP-A from CENP-B regions following auxin-induced degradation of CENP-C (Supplementary Fig. 5C-D), but we have not been able to statistically establish this effect.

### CENP-T intensity is decreased with auxin degradation of CENP-C and N

To verify that other CCAN proteins were affected by the loss of CENP-C and CENP-N, we measured the fluorescence of CENP-T labeled with fluorescent antibodies after CENP-C and CENP-N degradation, either on their own or simultaneously (Klare *et al*., 2015; Hoffmann *et al*., 2016). We used bundles of chromosomes (essentially the entire mitotic apparatus pulled from cells) to maximize fluorescence intensity and visualize a greater number of centromeres per cell. After degradation of CENP-C, CENP-N, and CENP-C/N (4- and 24-hour treatment), we continued to observe punctate dots of CENP-T, but at reduced intensity (approximately 30% of untreated) indicating that degradation led to appreciable loss of CENP-T from the CCAN, all of which were statistically significantly lower than the CENP-T signals from untreated cells (Fig. 6A), in accord with prior observations for CENP-C removal (Klare *et al*., 2015; Hoffmann *et al*., 2016). After quantification of the fluorescent intensity of the sprayed CENP-T, we noticed that there was no statistically significant difference in the fluorescent intensity between 4- and 24-hour auxin treatment in the CENP-C/N cell line or in the individual CENP-C or CENP-N lines (Fig. 6B). This suggests that the CCAN is rapidly compromised by degradation of CENP-C CENP-N, or both. Single data points for the fluorescence loss of CENP-T after CENP-C and N degradation in the chromosomal bundles can be seen in Supplementary Figure 6A.

**Figure 6.**
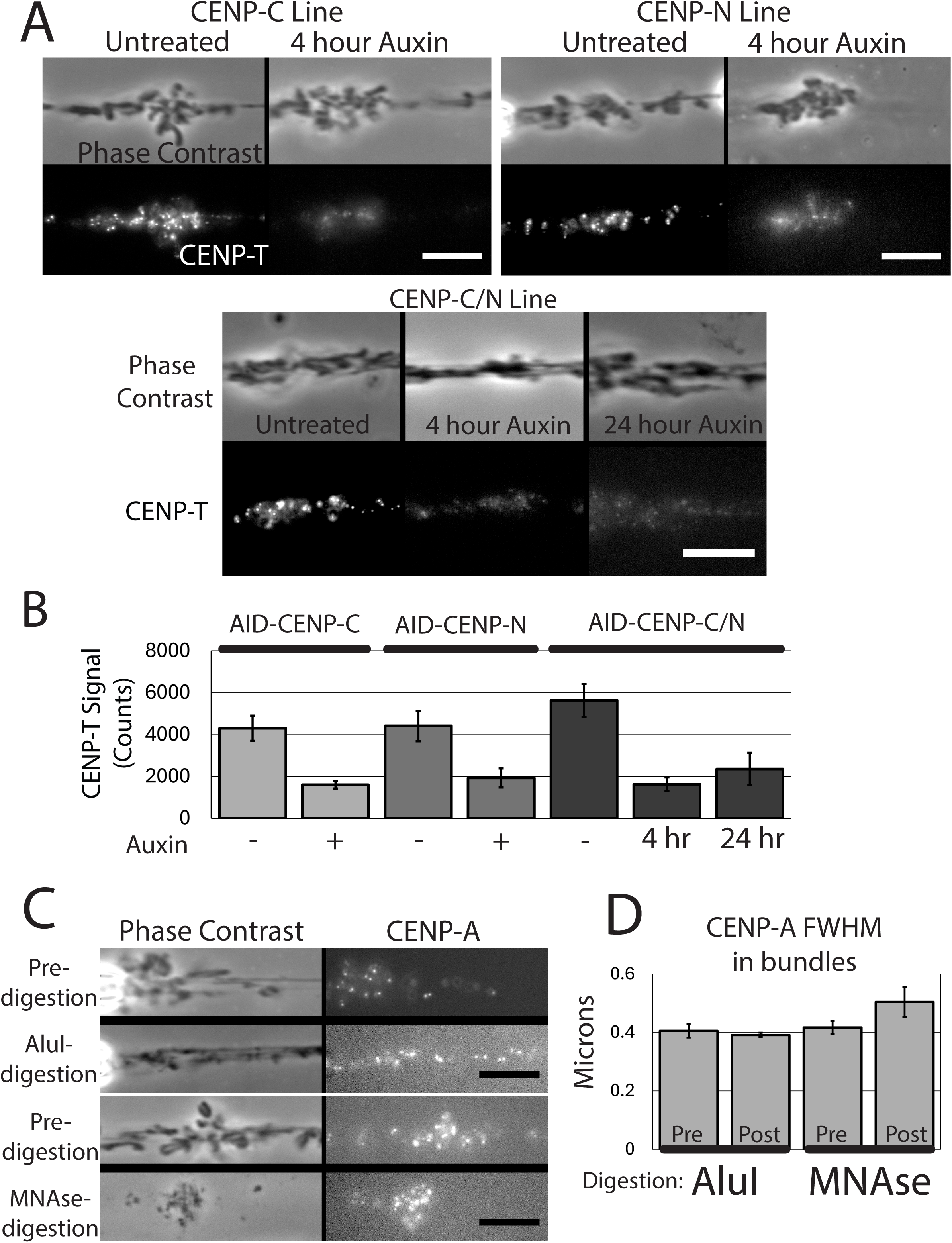
CENP-C and N degradation affects CENP-T protein levels while DNA digestion does not affect centromere length. (A) Fluorescence of bundles after CENP-T fluorescent antibody spraying in all conditions. The whole chromosome bundle in phase-contrast imaging is shown above each image of the fluorescent anti-CENP-T fluorescent antibody in its corresponding fluorescent channel across CENP-C, N, and C/N lines across untreated, 4-hour auxin-treated cells, and 24-hour auxin-treated cells in the CENP-C/N line. Scale bars 10 µm. (B)CENP-C, N, and C/N degradation over 4 and 24 hours decreases CENP-T fluorescence. Chromosome bundles were imaged in the anti-CENP-T fluorescence channel, where 5 random and in focus centromeres had their intensities measured at 500ms exposure time. In untreated cells, the bundles were bleached of fluorescence in the CENP-T fluorescent channel so there was no interference with the CENP-T fluorescence. In the CENP-C line bundles, the CENP-T fluorescence was 4300+/-600 counts above background (N=5) in untreated cells 1600+/-180 counts above background (N=3) (statistically significantly lower) in auxin-treated cells. In the CENP-N line bundles, the CENP-T fluorescence was 4410+/-730 counts above background (N=6) in untreated cells and 1930+/-460 counts above background (N=6) (statistically significantly lower) in auxin-treated cells. In the CENP-C/N line bundles, the CENP-T fluorescence was 5640+/-780 counts above background (N=4) in untreated cells, 1620+/-330 counts above background (N=3) (statistically significantly lower than untreated cells) in 4-hour auxin-treated cells, and 2360+/-780 counts above background (N=4) (statistically significantly lower than untreated cells) in 24-hour auxin-treated cells. Single data points are shown in Supplementary Figure 6A. (C) Example images pre- and post- AluI and MNase digestion on CENP-A length. AluI can digest isolated chromosomes and snap chromosome bundles (breaks AG^CT DNA sequences, which is also contained in human centromere repeat sequences). The isolated bundles were shown in both phase-contrast imaging and the CENP-A fluorescent channel, then sprayed with AluI mix and imaged again in phase-contrast imaging and the CENP-A fluorescent channel. An example of both pre and post digestion are shown in this image. MNase treatment is shown in the bottom panels and causes a near complete digestion of the chromosome except for the CENP-A region in phase-contrast. Scale bar 10 µm, lower right. (D)The centromere width does not change with AluI digestion. Bundles of chromosomes were extracted from the CENP-C line for use in digestion experiments, due to its higher intensity and endogenous CENP-A fluorescence. Before digestion, the bundles were imaged in phase contrast and the CENP-A fluorescent channel, sprayed with AluI or MNase, then imaged again in phase contrast and the CENP-A fluorescent channel. 5 random, in-focus centromeres were scanned with a boxplot (Fig. 1E,F) in both the pre and post digestion images, then averaged for the experimental data point. In these experiments, the CENP-A width was 0.41+/-0.02 µm (N=3) in undigested bundles, and 0.39+/-0.01 µm (N=3) (statistically insignificantly different) following AluI digestion. The CENP-A width was 0.42+/-0.02 µm (N=3) in undigested bundles and 0.49+/-0.03 µm (N=3) (statistically insignificantly different) in post MNase-treated bundles (there was more interference with the background in the post-MNase digested bundles due to the thorough digestion by MNase, complicating this result). Single data points are shown in Supplementary Figure 6B.

### CENP-A initial length is unchanged after digestion by a restriction enzyme and nuclease

We applied a more aggressive structural perturbation using DNA digestion to determine the impact on the CENP-A region. To do this, we compared the length of the endogenous CENP-A in the CENP-C line before and after a bundle was treated with AluI or MNase. AluI has previously been shown to digest chromosomes and cut chromosome bundles (Poirier and Marko, 2002; Sun, Kawamura, and Marko, 2011; Biggs *et al*., 2020) (see, *e.g.*, Genbank X02953.1, Genbank AF153368.2, Genbank M13882.1, Genbank X03113.1), while MNase can fully dissolve bundles. AluI cuts a sequence (AGåCT) found in the centromere satellite DNA in addition to many sites through the chromosome (Meneveri, Agresti, and Ginelli, 1984; Mezzanotte *et al*., 1992; Calabrese *et al*., 1994; Nieddu *et al*., 2003) and MNase cuts all free base pairs, which we reasoned might destabilize the centromere. The example image of the chromosome bundle in phase-contrast imaging and CENP-A fluorescence before and after AluI treatment is shown in Figure 6C (top) and MNase treatment (bottom). Comparing the average of five centromeres from each bundle, we found no statistically significant difference in the length of CENP-A signal before or after AluI or MNase digestion (Fig. 6D), indicative of a remarkable degree of stability. Single data points for the CENP-A average length in the bundles before and after digestion can be seen in Supplementary Figure 6B. (We attribute the post-MNase increase and spread in CENP-A lengths to the difficulty in isolating a clean CENP-A signal from the chromosome bundle dissolving away.) This is indicative of independence of the CENP-A puncta not being dependent on contiguity of accessible DNA for their structural maintenance.

## Discussion

### The core CENP-A centromeric region is not stretched by overall chromosome stretching

We studied the mechanical properties and behavior of the CENP-A-chromatin region of the human mitotic chromosome, to understand its structure and how it resists applied tension. We found that the centromere resisted all tension applied by stretching the chromosome along its long axis, such that even when the whole chromosome was increased in length by 300-700% (Fig. 3B, D-F of an example stretch) the centromere was essentially unstretched (Fig. 4B, D).

This striking result could be explained in two ways. First, the CENP-A region may be extremely stiff compared to the chromosome arms. In the extreme case where we imagine that all the tension in the chromosome is proportionally conducted through the CENP-A region, our results would indicate that the CENP-A region should have a doubling force at least 28 times larger than the chromosome arm as seen in the CENP-C line (the average CENP-A region stretches by at most 0.035 = 1/28 on average relative to the chromosome arms) (Fig. 4B, CENP-C - Auxin). Mitotic chromosome arms have a modulus of roughly 400-700 Pa (Supplementary Fig. 2B), indicating a modulus of at least 11-20 kPa for the CENP-A domain. This is a reasonable estimate for a protein-based structure, given that solidly bonded protein structures like f-actin and microtubules have Young’s moduli of about a GPa (Kojima, Ishijima, and Yanagida, 1994; Gardel *et al*., 2008). However, the result is still surprising, given that we were unable to observe any stretching of the chromatin containing CENP-A, while we observed the CENP-B-containing region of chromatin stretch with force.

The second possibility is that not all the tension in the chromosome is conducted through the CENP-A region in our chromosome-stretching experiments, *i.e.*, the CENP-A region is mechanically “decoupled” from the chromosome. The simplest way to envision this is to suppose that the CENP-A structure is lateral to the underlying chromatin, consistent with several models of centromeric formation and microscopy studies indicative of lateral displacement of the centromere from the chromosome body (Earnshaw, Ratrie, and Stetten, 1989; Lawrimore and Bloom, 2019, 2022; Lawrimore *et al*., 2022; Di Tommaso *et al*., 2023). Our observations that the two CENP-A regions (from each chromatid) can be pulled apart while the intervening CENP-B chromatin stretches (Fig. 5C, Supplementary Fig. 5) are also consistent with this second possibility (*i.e.*, the CENP-B region carries the mechanical stress). The CENP-A region could still be much stiffer than the other chromatin, but having it displaced from the underlying chromatin would greatly reduce the fraction of the total tension conducted through it, leading to the little-to-no deformation seen in our stretching experiments.

The mechanical disconnection of the CENP-A region from the rest of the chromosome along its long axis would also explain how other groups have been able to see deformation of the centromere and specifically the CENP-A region upon loss of CENP-C under merotelic attachments (Cimini *et al*., 2001; Suzuki *et al*., 2014). Under normal spindle attachments the mitotic spindles could provide up to hundreds of pN of force (Kajtez *et al*., 2016; Ye, Cane, and Maresca, 2016; Maiato *et al*., 2017; Anjur-Dietrich, Kelleher, and Needleman, 2021). We typically applied higher forces to the whole chromosome along its long axis in our stretching experiments (stretching 3-7 times its initial length, which would be well over 100 pN, given their average doubling force (Supplementary Fig. 2B), which is past the point of steepening of chromosome stiffening (Supplementary Fig. 2A)). We hypothesize that our observation of no centromere stretching reflects the difference in the way force is applied to the centromere in our experiments (the long axis of the chromosome) compared to merotelic attachments (direct attachments). Our results indicate that the chromatin structure of the centromere does not deform under large amounts of longitudinal chromosome stress, even when lacking the CCAN. Additionally, since we can see the separation of sister chromatid’s centromeres, we can assume that the chromatin between sister centromeres is also less stiff than centromeric chromatin or is not mechanically anchored to centromeric chromatin.

We also found that since the CENP-B-containing region of the pericentromere deforms, it is still mechanically in line with the long axis of the chromosome but is stiffer than the whole chromosome. These estimates of force could also give us an additional floor of stiffness for CENP-A of 70-fold stiffer than the chromosome arms, as we could not see a consistent increase of 75 nm (1 pixel) on a 750 nm object (10%) (CENP-A) (Fig. 4 B, D) while being stretched 7-fold (7/0.1). This could be somewhat due to the lower proportion of CENP-A in the CENP-B-containing portion of the chromosome. Again, this 70-fold increase in stiffness is not surprising from a protein-stiffness perspective but is surprising from a chromatin-stretching perspective.

### The core CENP-B centromeric region is longer than the CENP-A regions, is stretched by overall chromosome stretching, and is stiffer than the chromosome arms

Our experiments visualizing the CENP-B-containing chromatin highlight the differences in the relative properties of the CENP-A and CENP-B regions (Fig. 5C). On average, the 1200- nm-long CENP-B regions (Fig. 5B) are approximately 60% larger (longer) than the 700 nm-long CENP-A regions (Fig. 4B) for unstretched chromosomes, suggesting that the functional centromere (CENP-A occupation) is restricted to an appreciably smaller genomic region than that occupied by CENP-B, seen in other research (Kyriacou and Heun, 2018; Altemose *et al*., 2022; Chardon *et al*., 2022). When a chromosome is stretched, the CENP-B region stretches at a rate of about 35% that of the whole chromosome. This indicates that the CENP-B region has a modulus roughly three times that of the metaphase chromosome, or approximately 1400 to 2450 Pa, given the same cross-sectional area, which we typically observe after imaging fluorescent CENP-B.

In our imaging, CENP-B occupies the whole chromosome cross-section both before and during chromosome stretching. We often (about half of the time) observe the two CENP-A-containing/core-centromere regions move apart from one another (Fig. 5C, Supplementary Fig. 5). We also observe the two CENP-A regions to be displaced laterally from the more central CENP-B region. This supports the second possibility mentioned above, that not all (or perhaps little) of the longitudinal stress placed on the chromosome in our experiments goes through the CENP-A region, and instead passes through the CENP-B “core” (and possibly the Chromosome Passenger Complex or CPC, found between the centromeres and sister chromatids). This is consistent with prior observations of the CENP-B region connecting or lying underneath the CENP-A regions (Fachinetti *et al*., 2015; Fujita *et al*., 2015; Ohzeki, Otake, and Masumoto, 2020; Chardon *et al*., 2022).

### Degradation of CENP-N and CENP-C, either individually or simultaneously, does not alter whole chromosome, CENP-A, or CENP-B region stretching

We depleted the CCAN components CENP-C and CENP-N using an AID degron system, both individually, and simultaneously. We found that 90+% depletion of these proteins (estimated via remaining fluorescence, Fig. 2C, D) led to no change in the degree of stretching of the CENP-A region relative to the whole chromosome (Fig. 4B, D). However, the initial size of CENP-A shrunk following CENP-C/N, suggesting their depletion causes destabilization of some CENP-A, but the destabilized region is not mechanically affected (Fig. 4B, D). The continued mechanical stiffness of the centromere after CCAN disruption indicates that the CENP-A region itself (not the CCAN) can resist longitudinal forces without either CENP-C and CENP-N or that a very small residual population of CENP-N/C is sufficient to maintain the CENP-A region’s chromatin structure under mechanical stress. We also observed that none of the three degradation experiments (C, N and C+N) lead to any significant change in the untreated whole-chromosome stiffness (Supplementary Fig. 2B), or in stiffness of the CENP-B region (Fig. 5B). This is consistent with the CENP-B’s or whole chromosome’s chromatin structure being not directly dependent on the CCAN. This opens the possibility that the underlying chromatin may be responsible for the centromere region’s stiffness.

We note that our experiments with coincident knockdown of CENP-C and -N used antibody labeling of CENP-A and therefore could be affected by antibody crosslinking however this cannot explain the lack of stretching in individual CENP-C or CENP-N depletion experiments.

### Degradation of CENP-N or CENP-C leads to a 70% removal of CENP-T from centromeres

Antibody labeling of the centromeric histone analog CENP-T revealed that it was reduced by roughly 70% after degradation of either CENP-N, CENP-C, or CENP-N and -C together, in agreement with previous experiments (Fig. 6A, B) (Klare *et al*., 2015; Hoffmann *et al*., 2016). Disruption of the CCAN therefore destabilizes other CCAN components (Klare *et al*., 2015; Hoffmann *et al*., 2016); however, because CENP-T is a DNA-binding protein, it is reasonable that some of it should remain on DNA after CCAN component depletion. We found that simultaneous degradation of both CENP-C and CENP-N does not seem to have an additive effect in destabilizing the CCAN via CENP-T signal reduction, nor does long-term, 24-hour simultaneous degradation of both molecules have a more significant effect on CENP-T signal reduction in contrast to the other experiments.

### The CENP-A region is unaltered by digestion of Alu DNA

Prior experiments on whole chromosomes have shown that digestion of DNA by blunt- cutting restriction enzymes including by the 4-base-cutter AluI leads to dissolution of chromosomes, due to cutting of DNA between chromatin-crosslinking elements (Poirier and Marko, 2002; Sun, Kawamura, and Marko, 2011), while MNase dissolves chromosomes. The AluI sequence is also contained within human a-satellite DNA sequences, although the frequency of the sequence is higher in the chromosome arms. Digestion of bundles of chromosomes by AluI and MNase led to gradual dissolution of the chromosomes (as expected) but did not change the size of the CENP-A regions (Fig. 6C, D). This result agrees with studies of AluI cleavage protection in the centromeric regions of whole chromosomes, although note AluI sites are less prevalent in the centromere compared to the whole chromosome (Meneveri, Agresti, and Ginelli, 1984; Mezzanotte *et al*., 1992; Calabrese *et al*., 1994; Nieddu *et al*., 2003). All these results suggest that the CENP-A region structure, once formed, is not strongly dependent on DNA continuity, and is instead more dependent on other interactions, *e.g.*, protein-protein contacts or possible self-interactions.

### The CENP-A core of centromeres behaves as a protein-stabilized structure underlying the CENP-B chromatin

Our results indicate that the CENP-A-containing/core-centromeric region of the chromosome is a resilient nucleoprotein structure. We find that stretching the whole chromosome longitudinally did not deform CENP-A regions but deformed the CENP-B region. The CENP-B region appears to comprise the chromatin underlying the CENP-A regions, suggesting that the CENP-A regions are adjacent to, lateral to, or on top of CENP-B, rather than being in-line with the CENP-B region. Rather remarkably, nearly complete depletion of either or both of CENP-C and CENP-N did not alter the mechanical responses of the CENP-A or CENP-B regions. We believe that the CENP degradations may have destabilized the CENP-A region of the centromere, but the remaining region is still incredibly stiff or mechanically disconnected from the mechanical axis of the whole chromosome.

Our results on the relative location of CENP-A and CENP-B are consistent with recent ultrastructural studies (Di Tommaso *et al*., 2023) which observe that the α-satellite DNA sequences (probed directly using a hybridization method) are organized into a ring-like structure, which may be the result of looping of the DNA into a rosette (Fig. 5 of (Di Tommaso *et al*., 2023)), possibly by the action of SMC complexes at the base of the loops (Lawrimore and Bloom, 2019, 2022; Lawrimore *et al*., 2022). The actual images of the α-satellite DNA in that study are displaced from one another, with other DNA between them (Fig. 2, 3 of (Di Tommaso *et al*., 2023)), consistent with our observation of the CENP-A regions displaced laterally from one another by CENP-B-containing chromatin. This would also be consistent with the mechanically decoupled model of the CENP-A-containing region of the centromere and the central/longitudinal axis of the whole chromosome.

Our results are also consistent with models depicting the centromere being biased towards clustering together at the most lateral regions at the centromere (Lawrimore and Bloom, 2019, 2022; Lawrimore *et al*., 2022). These regions are also adjacent to the heterochromatin and SMC-complex-rich pericentromere that are distinct from the CCAN-binding region of the centromere. This model fits with our results as two (one per chromatid) distinct CENP-A regions can separate from each other, neither of which stretch under longitudinal, chromosome strain. Meanwhile, the underlying pericentromere, enriched in stiffening elements, carries the bulk of the stretching tension of the longitudinal, chromosome strain, but resists the pulling forces more compared to the chromosome arms. To test the idea that the CENP-A region flexibility might be impacted by removal of condensin (Ribeiro *et al*., 2009; Ribeiro, Vagnarelli, and Earnshaw, 2014), we did preliminary experiments where we depleted SMC2 from CENP-C cells via siRNA (Sun *et al*., 2018) and stretched the resulting chromosomes (Supplementary Fig. 6C, D). We found the SMC2siRNA reduced SMC2 antibody-stained fluorescence, but did not affect the stretch ratio of the chromosome bundles before or after SMC2siRNA treatment (Supplementary Fig. 6E, F). Additionally, pooling the data together, we found that there was no correlation between SMC2 fluorescence and CENP-A stretch ratio (Supplementary Fig. 6G). The lack of any deformation of the CENP-A region after SMC2 siRNA may be due to incomplete knockdown (especially if SMC2 is preferentially assembled at the centromere), or it may reflect the difference in cell types (Ribeiro *et al*., 2009; Ribeiro, Vagnarelli, and Earnshaw, 2014).

Figure 7 shows a model consistent with our results as well as those of (Di Tommaso *et al*., 2023), where CENP-B-rich chromatin in the two adjacent chromatids has the two CENP-A-rich regions organized at the surface of the chromosome away from the CPC-containing inner centromere. A fraction of α-satellite repeats contains CENP-B B-box binding sites, so it may be that CENP-B is distributed into the CENP-A region. For the type of structure shown in Fig. 7, longitudinal stretching of the chromosome will result in stretching of the underlying and adjacent CENP-B chromatin, but with little deformation of the CENP-A-rich loops, especially if the rosette structure envisioned by (Di Tommaso *et al*., 2023) is connected into the remainder of the chromosome at closely adjacent points (as shown). An open question is why the CENP-A-rich region should form a “looped-out” rosette structure (Fig. 5 of (Di Tommaso *et al*., 2023)), but this could be the result of sufficiently strong interactions between CENP-A molecules, perhaps induced by CCAN components.

**Figure 7.**
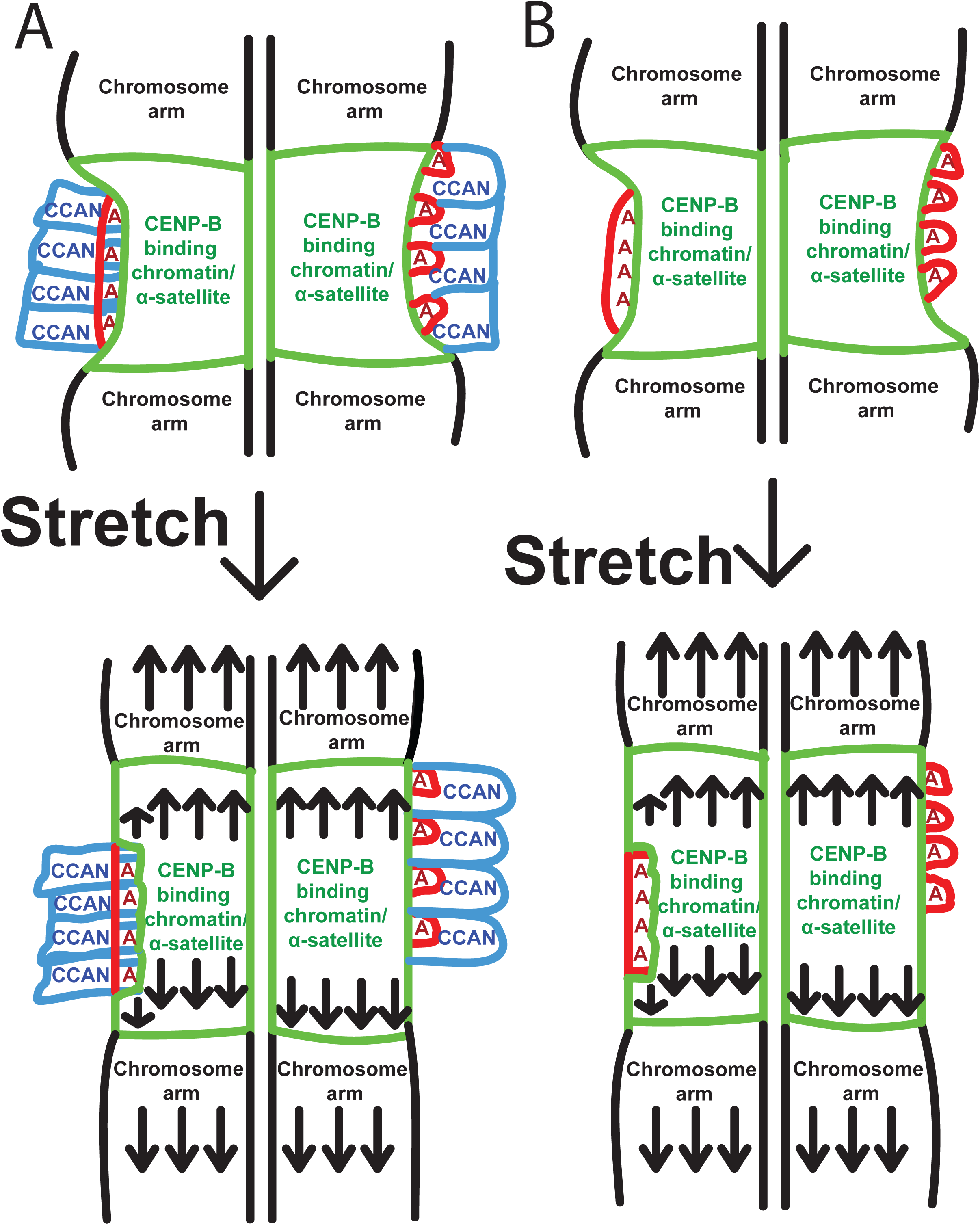
Model of chromosome stretching with centromere tracking and alternate possibilities. (A) Stretching schematic of the chromosome from untreated cells. The top panel shows a cartoon schematic of the unstretched chromosome arms colored in black. The chromosome arms are connected to the larger green CENP-B binding/α-satellite DNA. The red CENP-A nucleosomes are smaller and lateral to the CENP-B region. The CENP-A nucleosomes are connected to the blue CCAN structures that also invade the CENP-A domain, representing CENP-T binding to centromeric chromatin. The left side of the chromosome shows the model as if the CENP-A region is still along the chromosome axis and experiences stretching forces when the whole chromosome is stretched, while the right-hand side shows the model if the CENP-A region is mechanically disconnected from the central axis of the chromosome and thus does not experience forces when the whole chromosome is stretched. The bottom panel shows the example chromosomes under stress via stretching of the whole chromosome. The chromosome arms stretch the most, while the CENP-B region stretches less. This model shows the sliding of the two chromatid centromeres from each other in the model due to them not being locked into a specific connected region in the centromere relative to each other. Both panels show no change in the distance of the CENP-A region from the unstretched picture in the top panel. The left-hand panel shows how an increase in stiffness in the CENP-A region compared to the CENP-B region allows CENP-A to maintain its integrity despite being under stress, in that it resists the pulling tension more than the CENP-B region. The right-hand panel shows how if not in line with the chromosome-axis stretching, CENP-A would be under no/minimal stress, and thus not stretch. (B) Stretching schematic of chromosome from CCAN-degraded (auxin-treated) cells. The model shows the same effects of stretching as in (A), but without the CCAN. Since our experiments show that there is no change in stretching, nor in initial length, we show that the stretching effects are near-identical in the model but missing the structures in between the CENP- A nucleosomes.

Our result of invariance of the CENP-A mechanical response under CENP-N/C removal suggests that instead this strength might be due to the CENP-A chromatin itself. The stiffness in the chromatin itself could be facilitated by an increase in both heterochromatin and SMC complexes, like condensin and cohesin, localized around the centromere (Sun *et al*., 2018; Biggs *et al*., 2019; Strom *et al*., 2021). Further experiments directly deforming the centromere across its short axis, such as stretching the centromere by pulling it perpendicular to the chromosome axis (e.g., using manipulation via microneedle coated by antibodies to CENP-A), would be useful to further quantify centromere stability. The stress in our chromosome-stretching experiments might also be supported by the CPC, which connects the two centromeres. The CPC and CENP-B-rich regions are both candidate force-transmitting complexes that bypass the CENP-A region and would be interesting targets for future experiments of the type discussed in this work.

## Materials and Methods

### Cell culture, maintenance, and auxin treatment

All cells used in this study were modified DLD1 cells (human colorectal adenocarcinoma derived) containing an *Os*-TIR1 gene (E3 ubiquitin ligase targeting an auxin induced degradation (AID) tag upon auxin treatment) under a CMV promoter (continuously active). All cell lines used were previously studied and characterized in the 2018 Cao *et. al.* paper (Cao *et al*., 2018). Cell line 405 (CENP-C in the Figures and Results sections) contained a heterozygous C-terminal 3xFlag, Ruby fluorophore-containing CENP-A and a homozygous YFP fluorophore-containing, AID-tagged CENP-C. Cell line 408 (CENP-N in Figures 5, 6, and Supplementary Figures 1, 5, and 6) contained a heterozygous C-terminal 3xFlag, Ruby fluorophore-containing CENP-A and a homozygous sfGFP fluorophore-containing, AID-tagged CENP-N. Cell line 409 (CENP-N in Figures 2, 4, and supplementary Figures 1 and 4) contained a homozygous N-terminal 3xFlag, Ruby fluorophore-containing CENP-A protein and a homozygous sfGFP fluorophore-containing, AID-tagged CENP-N. Cell line 403 (CENP-C/N in the Figures and Results sections) contained a homozygous sfGFP fluorophore-containing, AID-tagged CENP-N and a homozygous Ruby fluorophore-containing, AID-tagged CENP-C.

All cells were maintained in RPMI1640 (Corning) complete media. Complete media contains an additional 10% fetal bovine serum (FBS) (HyClone) and 1% 100x penicillin/streptomycin (Corning) from the base media. The cells were incubated at 37°C and 5% CO_2_ for no more than 30 generations, passaged every 2-4 days. Experiments used cells that recovered 1-3 days after plating. Cells were freely cycling and not treated with drugs designed to synchronize the cells or otherwise affect the cell cycle. Auxin treatments were performed by aspirating the untreated media, then replacing it with the RPMI1640 complete media with 0.5 mM Indole-3-Acetic Acid (IAA) 4 hours before performing the experiment, unless specifically labeled as a 24-hour treatment (only in the C/N cell line), where the auxin-containing media replaced the untreated media 20-28 hours before performing the experiment.

### Single chromosome and bundle capture

All experiments used an inverted microscope (IX-70; Olympus) with a 60x 1.42 NA oil immersion objective with additional 1.5x magnification at room temperature and atmospheric CO_2_ levels. We performed experiments in under 3 hours after removal from the incubator to minimize damage to the cells being analyzed. Figure 3A depicts the steps to isolating a mitotic chromosome from a mitotic cell. Prometaphase cells were identified by eye (upper left) and lysed with a pipette (WPI TW100-6) containing 0.05% Triton-X 100 (US Biologicals) in PBS (Lonza) (upper middle). All non-lysing pipettes were filled with PBS. After lysis, the bundle of chromosomes would be stabilized with a pipette connected to a manual micromanipulator and moved away from the lysed cell (upper right). For bundle experiments, the bundle would be aspirated by two pipettes on opposite sides to stabilize it for spray experiments using MP285 micromanipulators (Sutter Instruments). In single chromosome experiments, one end of a random, loose chromosome was grabbed by the deflection pipette (WPI TW100F-6) using a MP285, moved from the bundle, and subsequently grabbed with the pulling pipette on the other end (lower left) with another MP285. The bundle was then removed to isolate the target chromosome (lower right).

### Live stretch-deflection tracking

Once captured, the attached pipettes were moved perpendicular to the chromosome, stretching the chromosome to roughly its native length. The pulling pipette was then commanded to move 20 µm and return to the starting position using a custom LabView program that interfaced with the MP285. The commanded stretch would typically result in an actual pull of 10-15 µm. The pulling pipette moved at a constant rate of 1.00 µm/sec (actual rates about half the commanded rate) in 0.2 µm steps using the LabVIEW program, while tracking the stiff and force pipettes. This experiment was performed before the multi-fold chromosome stretch used to obtain the centromere vs chromosome stretch data. Figure 3B shows an example of a stretched chromosome in discrete steps (stretched further than 20 µm for demonstration purposes). Figure 3C shows an example LabVIEW deflection vs stretch trace and a linear-regression-derived slope of the trace. The stretch- deflection slope was multiplied by the spring constant of the deflecting pipette to obtain the spring constant of the chromosome. Each chromosome was stretched at least 3 times and typically 6 times to provide an accurate and reproducible measurement of its spring constant with the coefficient of variation typically under 0.2. The deflecting pipette’s spring constant was found by measuring the ratio of its resistance to pushing a pipette of known stiffness to the known pipette’s full movement. The chromosome’s spring constant was multiplied by its initial length (the distance between the centers of the pipettes while the chromosome was under little to no force nor buckling) to obtain its doubling force. In all auxin treatments we found no change in their doubling force compared to the untreated chromosomes (Supplementary Fig. 2B, top). We also found no change in their elastic modulus (doubling force divided by the cross-sectional area of the chromosome) between untreated and auxin-treated chromosomes (Supplementary Fig. 2B, bottom). For all doubling force experiments, we stretched the chromosome in its linear, non-hysteresis regime of the chromosome, but in the centromere stretching experiments, we would easily reach the stretching regime where the chromosome experienced plastic deformation and a non-linear stretch- deflection slope, typically found above four-fold its initial length (Supplementary Fig. 2A).

### Measuring centromere and chromosome stretching for mitotic chromosomes

For centromere stretching experiments, we stretched the whole isolated chromosome a large amount (up to the diagonal of the whole screen at 90x magnification, typically ∼7 fold the chromosomes’ initial length) and imaged the chromosome in phase contrast (Fig. 3B) to obtain the chromosome length, then imaged the chromosome in the fluorescence channel for the centromere length (Fig. 3D). We wish to specifically mention that the amount the chromosome stretched was done via manual measurements using a centromere: chromosome length ratio, contrasted with the LabVIEW-based tracking software methods used in determining the whole-chromosome mechanics. This was repeated 3-15 times, depending on the stability of the fluorescence, resetting the chromosome to its initial length once the chromosome was stretched the full length of the screen. The fluorescent image of the centromere was used to trace its intensity (Fig. 3D row 4, E), which was fit with a gaussian curve. The length of the centromere was calculated as the full width at half-maximum (FWHM) of the gaussian fit. The centromere length was plotted against the chromosome length, both divided by initial conditions (Fig. 3F). The slope was derived by using a least-square fit linear regression to obtain the centromere: chromosome stretch ratio. Figure 3F shows an example trace of how CENP-A typically does not deform when the whole chromosome was stretched and how CENP-B typically stretched at 40% the rate of the whole chromosome.

### Chromosome bundle and isolated chromosome spray experiments

In spray experiments, the target object (isolated chromosome or chromosome bundle) was raised above the glass surface using the controlling pipettes. A wide-bore pipette filled with the spray liquid held in a perpendicular pipette holder would then be positioned close to the target object. The liquid would then be steadily sprayed onto the target by adjusting hydraulic pressure via liquid level of the water in the connected syringe. We used a target time of 5-15 minutes to dispense the 14 µL payload for the liquids. For the fluorescent measurements, the liquid mixture contained 2% primary or secondary antibody, 10% casein (Thermo scientific) (as a blocking reagent), 38% molecular biology grade water (Lonza), and 50% PBS. The fluorescent secondary antibody used was always an Alexafluor-488 conjugated goat anti-rabbit antibody (Invitrogen A11034). The primary antibodies were rabbit anti-target (CENP-A, B (Santa Cruz Biotechnology sc-376392), T, and SMC2 (Novus NB-100-373)). Rabbit polyclonal antibodies to CENP-A and CENP-T were generated in the Straight laboratory and affinity purified using the antigen coupled to solid support. The digestion experiments used 2% AluI or MNase (NEB), 10% NEB CutSmart buffer or MNase buffer, and 88% molecular biology grade water.

### Centromere size estimate correction using point-spread distance

Since we were using fluorescence as a readout for centromere lengths in the few hundred nm range and near our optical spatial resolution, we needed to determine the length resolution limitations of our microscopy setup. We did this by observing the point-spread function for a small, fluorescent source of known size. We used fluorescent spheres of known size 175 +/- 5 nm (Molecular probes P7220 component B) and the 1.0, 0.2, and 0.1 µm beads from the T14792 tetraspeck fluorescent microspheres kit (Invitrogen) adhered to a cover slip. We measured FWHM of their images (Supplementary Fig. 7A) and derived the point spread function as the square root of the measured FWHM squared minus the molecule size squared for each sized particle. We then averaged the numbers derived from these tests and found that our microscope had an average point spread function of 0.365 μm, which we used to correct the lengths of the CENP-A and CENP-B regions.

We used the measured average point source diameters to correct the lengths for CENP-A and CENP-B (Supplementary Fig. 7B) in the experiments, by taking the difference of the squares of the observed CENP-A/B length and the point-source diameter, and then taking the square root of that difference. This corresponds to a model of the CENP-A/B image being the convolution of a “true” image with a Gaussian point-spread function.

### Checking repeated exposure of centromeres to centromere length

We extracted single chromosomes from the CENP-C line and exposed the centromere 5 times to 5 seconds of fluorescent light (the length used to capture images of the centromere) to photobleach the centromere (Supplementary Fig. 7C). We then compared the FWHM of the centromere against the exposure number and found there was no consistent, nor large effect of exposure number on the measurement (Supplementary Fig. 7D). This finding led us to not change the length of the FWHM measurements with respect to exposure number.

### Fluorescent imaging of live cells, bundles, and isolated chromosomes

We imaged the fluorescent intensity of the endogenous fluorescent proteins in the cells before isolating a chromosome from a cell. The fluorescent signal of the centromere was calculated by drawing a 10-by-10 square around the centromere, using the CENP-A fluorescence for determining the location of the centromere, where able (Fig. 2A). The background fluorescence was taken from a centromere-free segment of the cell. We analyzed a population of cells with three independent runs untreated or with auxin treatment to obtain the expected distribution of cellular fluorescence of these cells, which were also treated with Hoechst (Life technologies). Hoechst was

not used in experiments with isolated chromosomes, so the mechanics of the chromosome were unaffected.

### SMC2 siRNA treatment

We removed the RMPI1640 complete media then treated 70% confluent cells with 100 nM SMC2 siRNA (Sun *et al*., 2018) in Optimem (Gibco) with 2% Lipofectamine 3000 (250uL Optimem, 5uL Lipofectamine 3000 (Invitrogen), and 2uL SMC2 siRNA 20μM stock (Sun *et al*., 2018)) for 6 hours. We then replaced the media with RPMI1640 complete and let the cells recover 16-40 hours and performed the bundle experiments on chromosome bundles from the cells.

## Statistics

For all our bar graphs and reported values, we display and state the average +\- the standard error of the mean in the bar graphs. We removed outlier data points via a rolling Grubbs outlier test. Statistical significance was determined by a student T-test, significance defined as falling under a P value of 0.05 and explicitly stated when between 0.05 and 0.10. All figures with single data points are included in the supplement with a spreadsheet of all data points. CENP-C data is shown in light grey bar graphs (Figures) and blue in individual data points (Supplementary Figures). CENP-N data is shown in medium grey bar graphs (Figures) and red, orange, and yellow in individual data points (Supplementary Figures). CENP-C/N data is shown in dark grey bar graphs (Figures) and purple or pink in individual data points (Supplementary Figures). Our N are in reference to different biological replicates per cell, per chromosome, or per bundle, except in reference to CENP-A, where N refers to the number of centromeres (periodically, our chromosomes would have two centromeres from each chromatid and counted where possible.)

## Acknowledgements

This work was supported by NIH grant R01-GM105847 (JFM), by subcontract to the University of Massachusetts under NIH grant UM1-HG011536 (Center for 3D Structure and Physics of the Genome, 4DN Consortium, JFM), and by NIH grant R01-GM074728 (AFS).

