## Supplementary Figures for "Independence of Centromeric and Pericentromeric Chromatin Stability on CCAN Components"

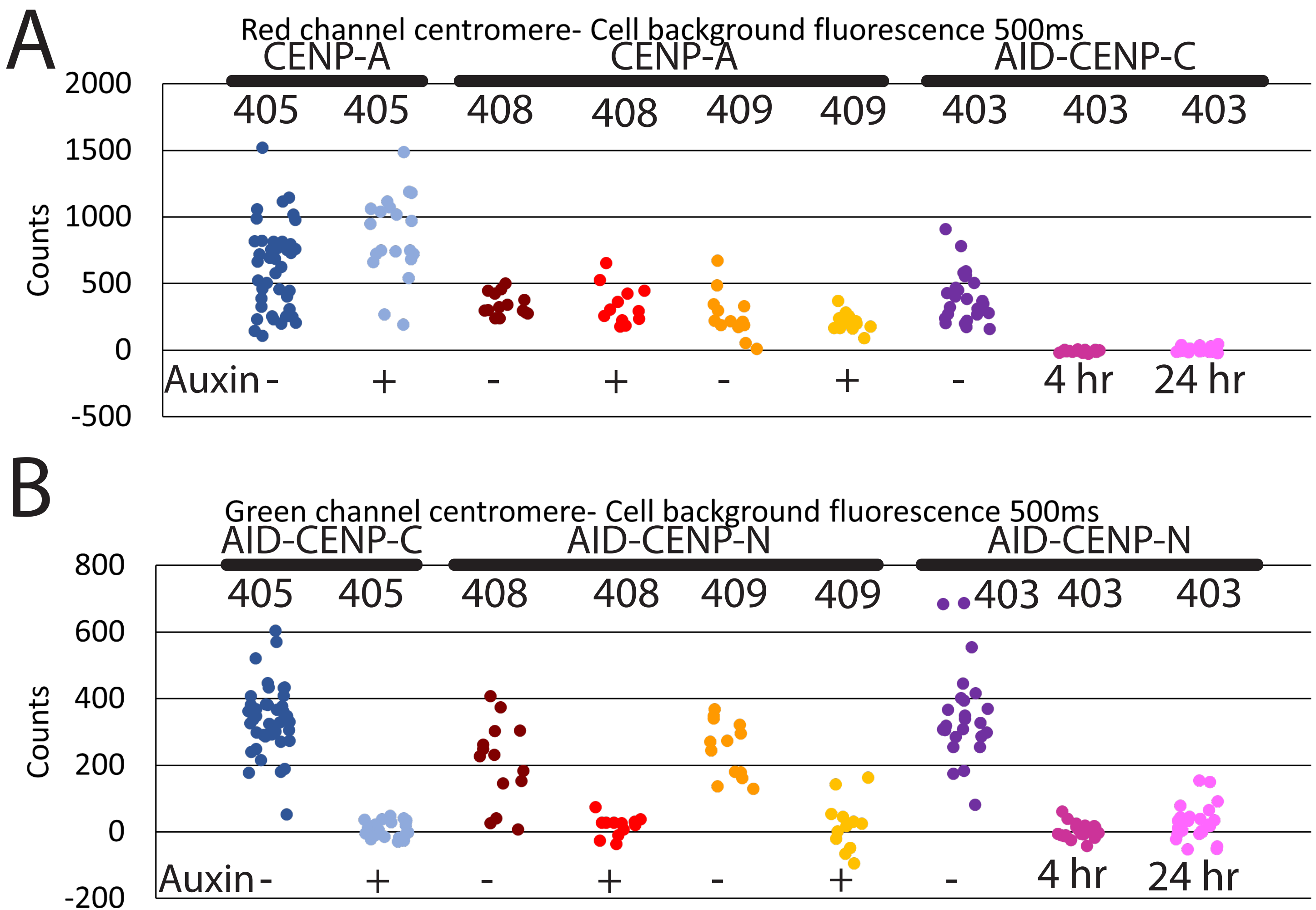

A
Red channel centromere- Cell background fluorescence 500ms
CENP-A
CENP-A
AID-CENP-C
405 405 408 408 409 409 403 403 403
Counts
2000
1500
1000
500
0
-500
Auxin - + - + - + - 4 hr 24 hr
B
Green channel centromere- Cell background fluorescence 500ms
AID-CENP-C
AID-CENP-N
AID-CENP-N
405 405 408 408 409 409 403 403 403
Counts
800
600
400
200
0
-200
Auxin - + - + - + - 4 hr 24 hr

Supplementary Figure 1. Single data points for the fluorescence intensity of centromere proteins in the cell.

(A) Single data points for the red channel fluorescence in each cell line before cell lysis. Averages, standard errors, and statistical differences of these lines can be seen in Figure 2C. Cell lines 405 (CENP-C) and 408 (CENP-N) contained a heterozygous C-terminal 3xFlag, Ruby fluorophore-containing CENP-A, which appeared in the red-fluorescent channel and did not degrade with auxin treatment, as expected. Cell line 408, like 409 did not show a statistically significant difference following auxin treatment in the red-fluorescent channel (343+/-22 (N=14)) counts above background in the untreated cells and 341+/-43 counts above background in cells following auxin treatment). Cell line 409 (CENP-N) contained a homozygous C-terminal 3xFlag, Ruby fluorophore-containing CENP-A, which appeared in the red-fluorescent channel and did not degrade with auxin treatment, as expected. Cell line 403 (CENP-C/N) contained a homozygous Ruby fluorophore-containing, AID-tagged CENP-C, which appeared in the red-fluorescent channel and degraded with both 4-hour and 24-hour auxin treatment, as expected. The 24 hour treated cells had an average fluorescent intensity of 4+/-4 counts above background (N=25), which was statistically significantly lower than the untreated cells.

(B) Single data points for the green channel fluorescence in each cell line before cell lysis. Averages, standard errors, and statistical differences of these lines can be seen in Figure 2D. Cell line 405 (CENP-C) contained a homozygous YFP fluorophore-containing, AID-tagged CENP-C, which appeared in the green-fluorescent channel and degraded with auxin treatment, as expected. Cell lines 408 and 409 (CENP-N) contained a homozygous sfGFP fluorophore-containing, AID-tagged CENP-N, which appeared in the green-fluorescent channel and degraded with auxin treatment, as expected. Cell line 408, like 409 showed a statistically significant decrease in intensity following auxin treatment in the green-fluorescent channel (203+/-8 (N=14) counts above background in the untreated cells and 17+/-9 counts above background in cells following auxin treatment). Cell line 403 (CENP-C/N) contained a homozygous sfGFP fluorophore-containing, AID-tagged CENP-N, which appeared in the red fluorescent channel and degraded with both 4-hour and 24-hour auxin treatment, as expected. The 24 hour treated cells had an average fluorescence intensity of 29+/-10 counts above background (N=25), which was statistically significantly lower than the untreated cells.

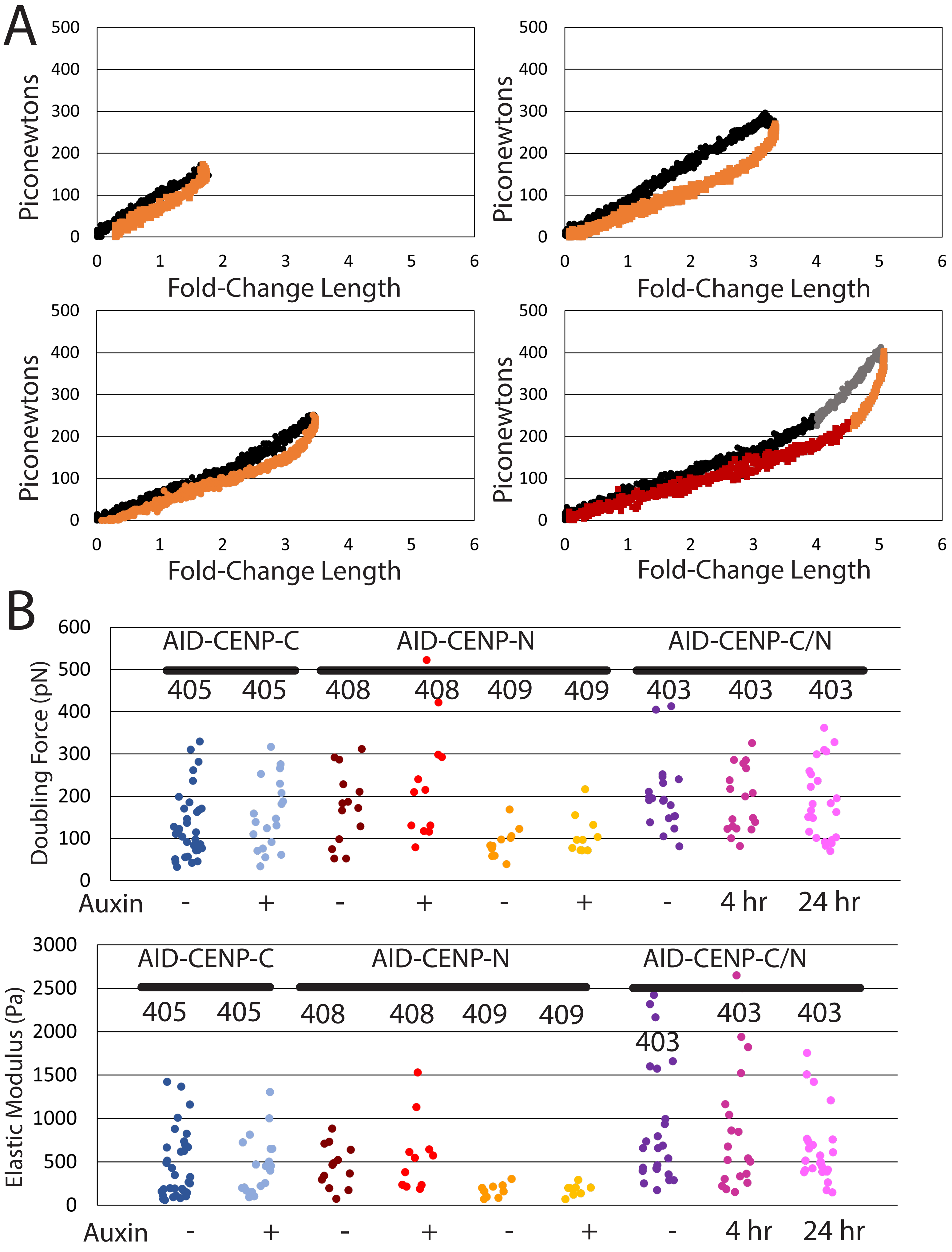

A
Piconewtons
Fold-Change Length
B
Doubling Force (pN)
AID-CENP-C
AID-CENP-N
AID-CENP-C/N
405
405
408
408
409
409
403
403
403
Auxin
-
+
-
+
-
+
-
4 hr
24 hr
Elastic Modulus (Pa)

Supplementary Figure 2. Chromosome cell line mechanics and large stretching effects.

(A) Example trace of a force vs chromosome extension for small and large stretches. The graphs show the outgoing curves in black or grey circles and a return curve in orange or red squares. The different graphs show the different behavior of an example chromosome under small or large stretching. The y-axis shows the deflection of the deflecting pipette multiplied by its spring constant (resulting in pN units), while the x-axis shows the amount the chromosome stretched (the pull pipette movement minus the deflection pipette movement) divided by the chromosome's initial length (resulting in a unitless measurement). The upper left shows the regime of stretching that is completely reversible and linear along the force-extension space. The upper right shows the regime where the stretch of the chromosome starts to show hysteresis (the return curve deviates from the outgoing curve) and weakens with additional stretches. The lower left shows the pull following the upper right's pull, which pulls the same distance as the upper right, but occurs under less force. This follow-up pull shows the chromosome is weakened after the second pull but does not show further weakening or hysteresis afterwards. The lower right shows the stretching length where pulling the chromosome more causes the force-extension slope to undergo a sharp increase in the force-extension curve (black to grey), meaning it becomes harder to stretch the chromosome. The return path of the lower right graph also shows that stretching the chromosome to such a degree (~4+ fold its initial length) results in a large amount of hysteresis. The chromosome undergoes a rapid decrease in force when shortening the chromosome, followed by an extended run with a smaller slope when returning to the 0-0 point in red.

(B) Auxin treatment causes no statistically significant change in chromosome doubling force. After chromosome isolation, we would stretch the chromosome and determine its doubling force as described in the Material and Methods. Each column is shown as a collection of individual data points. Untreated cells are shown in a darker color, while auxin treatments are shown in lighter colors. The CENP-C line is shown in dark and light blue, the CENP-N lines are shown in dark and bright red (408) or orange and yellow (409), and the CENP-C/N lines are shown in purple, magenta, and pink, becoming lighter with additional auxin treatment. The average doubling force of whole chromosomes in the CENP-C line was 131+/-142 pN (N=34) in untreated cells and 156+/-48 pN (N=20) (statistically insignificantly different) in auxin-treated cells. The average doubling force of whole chromosomes in the 408 CENP-N line was 175+/-23 pN (N=14) in untreated cells and 232+/-39 pN (N=12) (statistically insignificantly different) in auxin-treated cells. The average doubling force of whole chromosomes in the 409 CENP-N line was 91+/-11 pN (N=11) in untreated cells and 110+/-15 pN (N=10) (statistically insignificantly different) in auxin-treated cells. The average elastic modulus of whole chromosomes in the CENP-C/N line was 202+/-18 pN (N=21) in untreated cells, 190+/-18 pN (N=18) (statistically insignificantly different from all treatments) in 4-hour auxin-treated cells, and 188+/-18 pN (N=24) (statistically insignificantly different from all treatments) in 24-hour auxin-treated cells. The average elastic modulus of whole chromosomes in the CENP-C line was 452+/-66 Pa (N=34) in untreated cells and 465+/-76 Pa (N=19) (statistically insignificantly different) in auxin-treated cells. The average elastic modulus of whole chromosomes in the 408 CENP-N line was 454+/-67 Pa (N=13) in untreated cells and 573+/-127 Pa (N=11) (statistically insignificantly different) in auxin-treated cells. The elastic modulus force of whole chromosomes in the 409 CENP-N line was 170+/-23 Pa (N=10) in untreated cells and 182+/-21 pN (N=9) (statistically insignificantly different) in auxin-treated cells. The average elastic modulus of whole chromosomes in the CENP-C/N line was 886+/-141 Pa (N=24) in untreated cells, 922+/-173 Pa (N=20) (statistically insignificantly different from all treatments) in 4-hour auxin-treated cells, and 652+/-85 Pa (N=24) (statistically insignificantly different from all treatments) in 24-hour auxin-treated cells.

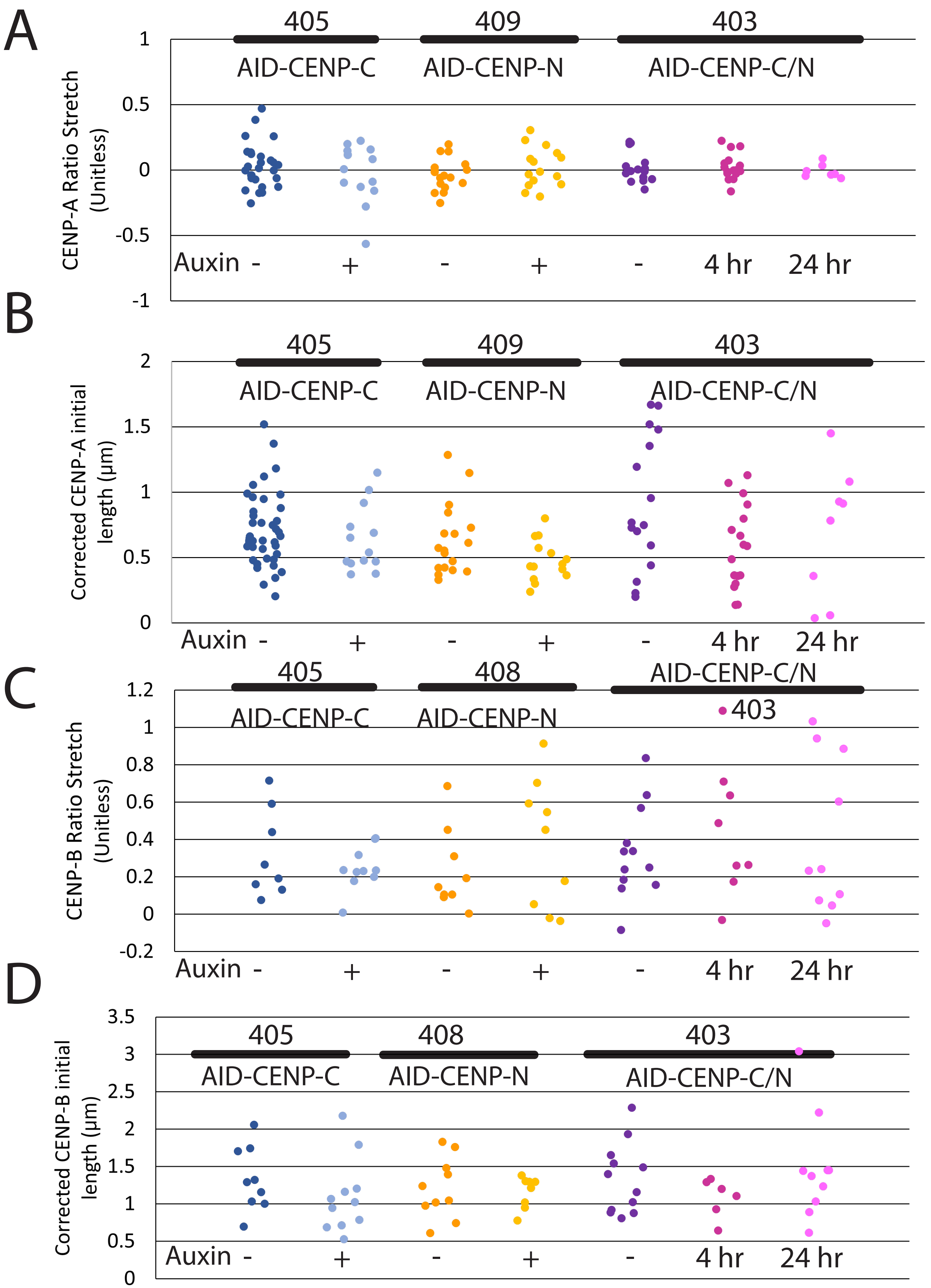
A
405
409
403
AID-CENP-C
AID-CENP-N
AID-CENP-C/N
CENP-A Ratio Stretch (Unitless)
1
0.5
0
-0.5
-1
Auxin -
+
-
+
-
4 hr
24 hr
B
405
409
403
AID-CENP-C
AID-CENP-N
AID-CENP-C/N
Corrected CENP-A initial length (µm)
2
1.5
1
0.5
0
Auxin -
+
-
+
-
4 hr
24 hr
C
405
408
AID-CENP-C/N
403
AID-CENP-C
AID-CENP-N
CENP-B Ratio Stretch (Unitless)
1.2
1
0.8
0.6
0.4
0.2
0
-0.2
Auxin -
+
-
+
-
4 hr
24 hr
D
405
408
403
AID-CENP-C
AID-CENP-N
AID-CENP-C/N
Corrected CENP-B initial length (µm)
3.5
3
2.5
2
1.5
1
0.5
0
Auxin -
+
-
+
-
4 hr
24 hr

Supplementary Figure 3. Single Data Points for Centromeric Stretching.

(A) Single Data Points for CENP-A stretch ratio. The average graphs from Figure 4B and 4D, left hand side shown in the single data point spread instead of average with error bars. In short, all cell lines showed no statistically significant change in the CENP-A stretch: chromosome stretch following auxin treatment.

(B) Single Data Points for CENP-A initial length. The average graphs from Figure 4B and 4D, right hand side shown in the single data point spread instead of average with error bars. In short, there was no statistically significant change in CENP-A initial length in the CENP-C line following auxin treatment, a near statistically significant decrease in CENP-A initial length in the CENP-N line following auxin treatment, a statistically significant decrease in CENP-A initial length in the CENP-C/N line following auxin treatment for 4 hours, but no statistically significant difference in the initial length of CENP-A following auxin treatment for 24 hours to the untreated or 4 hour treated centromeres.

(C) Single Data Points for CENP-B stretch ratio. The average graphs from Figure 5B, top shown in the single data point spread instead of average with error bars. In short, all cell lines showed no statistically significant change in the CENP-B stretch: chromosome stretch following auxin treatment.

(D) Single Data Points for CENP-B initial length. The average graphs from Figure 5B, bottom shown in the single data point spread instead of average with error bars. In short, all cell lines showed no statistically significant change in the initial length of CENP-B following auxin treatment.

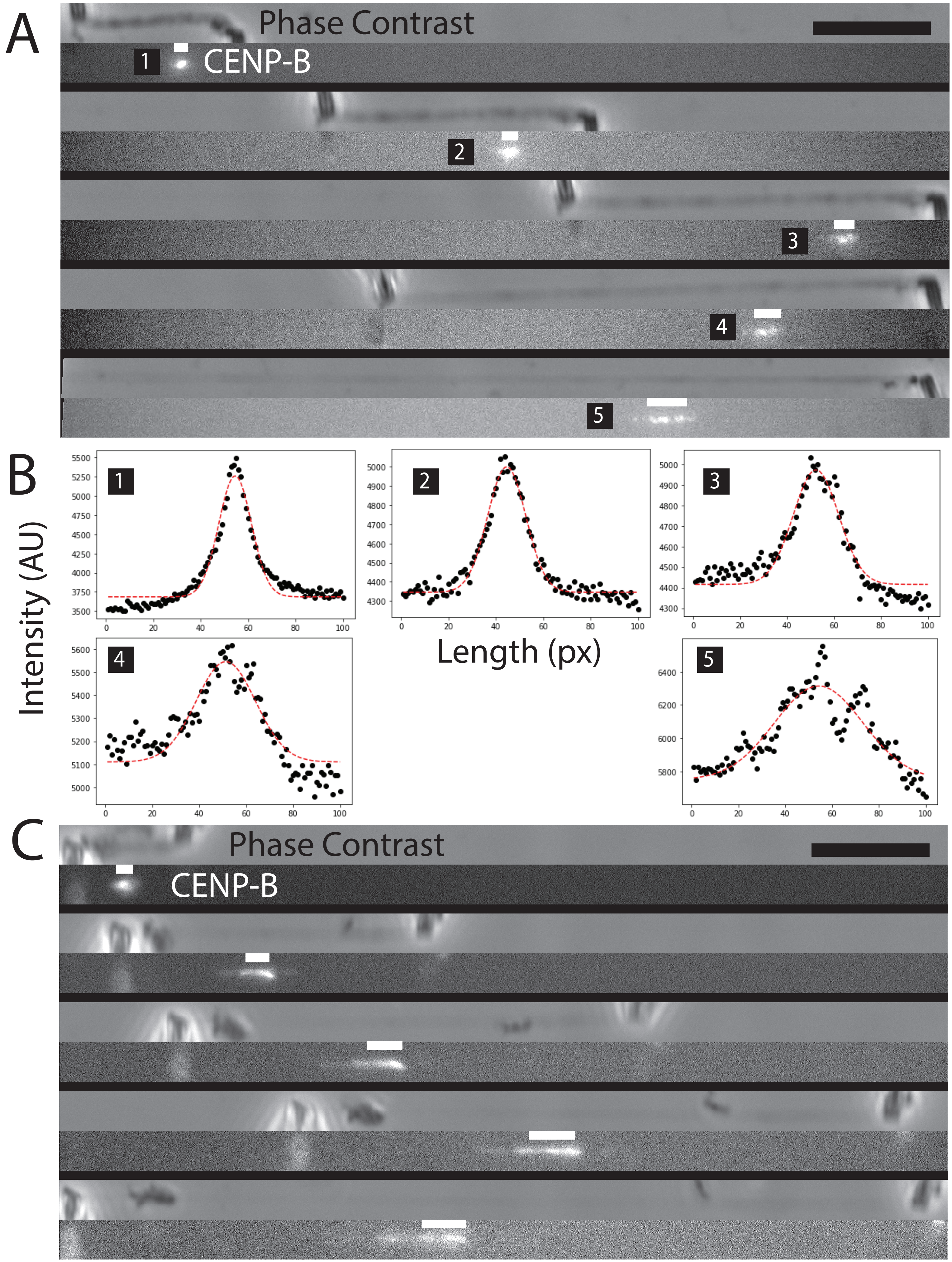
A
Phase Contrast
CENP-B
1
2
3
4
5
B
Intensity (AU)
Length (px)
C
Phase Contrast
CENP-B

Supplementary Figure 4. Example CENP-B stretching and example traces.

(A) Image of CENP-B shattering and stretching with chromosome stretching. Representative image of the antibody-stained CENP-B splitting into distinct punctate particles. The image shows a CENP-B stretch ratio of 0.338 and a corrected initial length of 1.16 μm. The FWHM program/script calculated size of CENP-B is shown in a white rectangular bar above the fluorescent signal, while the specific traces are shown in (B), each subpanel number corresponds with the number in (B). Scale bar (black) 10 µm long, upper right.

(B) Traces of the CENP-B signal from each panel in (A). Each plot represents the average box plot of the fluorescent signal in (A) with the number in the upper left-hand corner corresponding to the number in the left-hand side of the fluorescent channel in (A). The Y-axis shows the average intensity along the line in pixels (shown as the X-axis). The fracturing of the centromere can be seen in distinct but overlapping peaks of the fluorescent trace. Even though the peaks are spread along the fluorescence trace, the methods of fitting the curve to a gaussian fit still gives an acceptable estimation for the length of the centromere. This typically reveals a linear relationship between the chromosome stretch and the centromeric stretch as shown in Figure 3F.

(C) Image of CENP-B smearing and deforming with chromosome stretching. A repeat description of A, but with a centromere that stretches via smearing instead of cracking/fracturing with no corresponding traces. The image shows a CENP-B stretch ratio of 0.638 and a corrected initial length of 1.29 μm. White images above centromere show program-based FWHM-calculated length of the signal, Scale bar (black) 10 µm long, upper right.

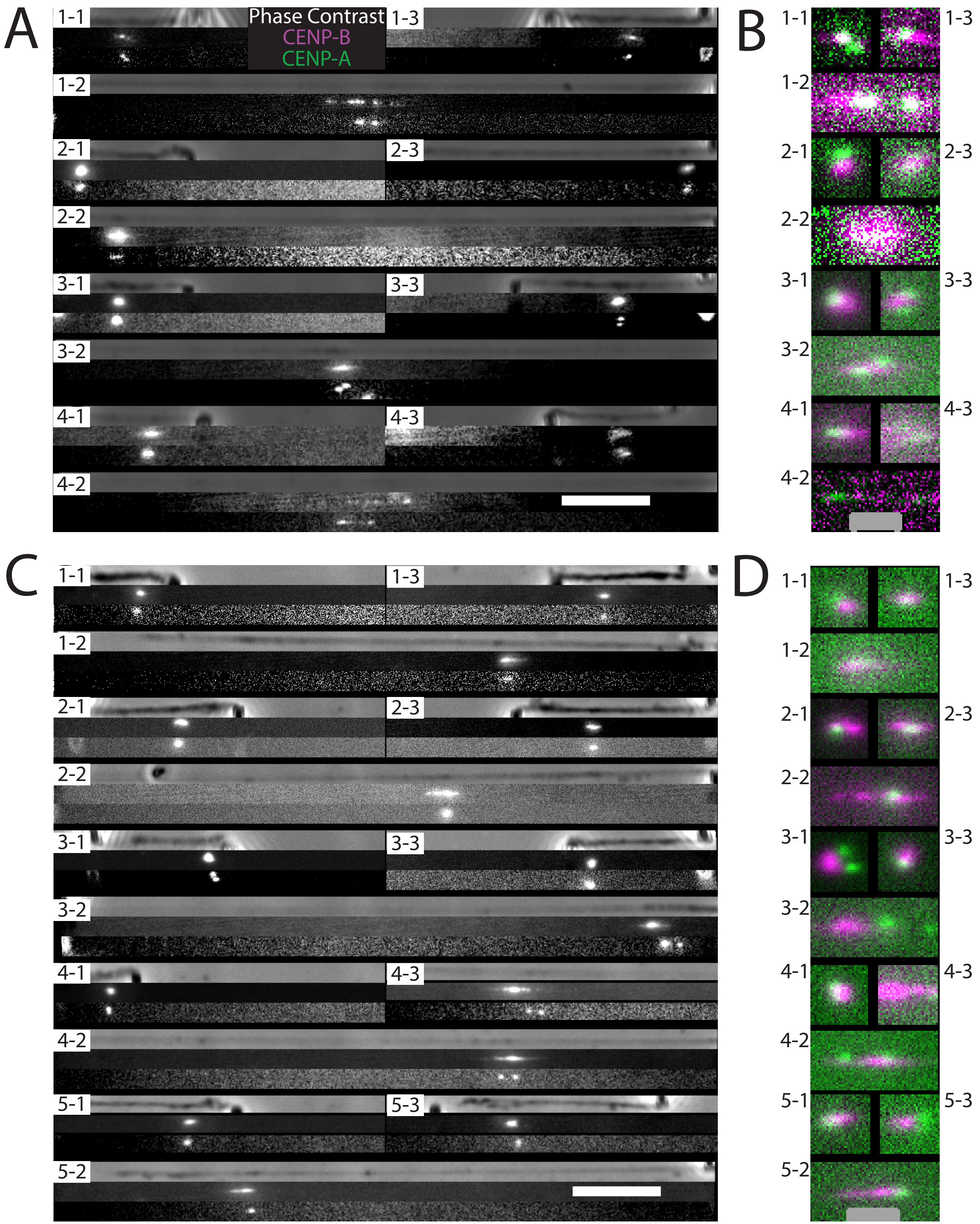
A
B
C
D
Phase Contrast
CENP-B
CENP-A
1-1
1-2
1-3
2-1
2-2
2-3
3-1
3-2
3-3
4-1
4-2
4-3
5-1
5-2
5-3

Supplementary Figure 5. Additional CENP-A and CENP-B costaining.

(A) CENP-A does not stretch while CENP-B stretches when the chromosome is stretched. We took a coincident picture of a 488-nm-conjugated antibody against CENP-B in the 405 cell line, which has a different colored (Ruby) CENP-A. 4 example chromosomes (4-#) are shown under initial conditions (top left panels), in stretched conditions (bottom panels), and in return conditions (top right panels). These panels are also labeled with the chromosome number-the chronological step in the stretching (upper left unstretched (#-1), bottom fully stretched (#-2), and upper right return to initial length (#-3). The phase-contrast imaging at the top of each set is used to show the whole chromosome, the CENP-B signal in the middle of each set is used to show its fluorescence and relative position, and the CENP-A signal at the bottom of each set is used to show its fluorescence and relative position. The nine images are repeated four times for four untreated chromosomes. The CENP-A from chromosomes 2, 3, and 4 can be seen separating from their sister chromosomes. Scale bar 10 µm long, bottom right.

(B) Colocalized signal of CENP-A and CENP-B zoomed in at the centromere. Each image overlaps the signals of the CENP-A (green) and CENP-B (magenta) signals of the images to the left, at the centromere, at 3x its original sizing to demonstrate the overlap or comparative location of CENP-A and CENP-B. Unstretched chromosomes shown in the top panels and stretched chromosomes shown in bottom panels similar to the numbering scheme in (A). Scale bar 2 µm long, bottom, in grey.

(C) Auxin treatment of the CENP-C cell line shows similar CENP-A and CENP-B stiffness as compared to untreated chromosomes. The same procedure as in (A) was repeated for the 5 chromosomes from auxin-treated cells. We again used the repetition of the three chromosome lengths in the initial, return, and stretched imaged in the phase-contrast, CENP-A, and CENP-B channels. The CENP-A from chromosomes 2, 3, and 4 can be seen separating from their sister chromosomes. The pictures show CENP-A and CENP-B may become somewhat decoupled (higher distance) following CENP-C degradation and chromosome stretching. Scale bar 10 μm long, bottom right.

(D) CENP-C line zoomed in and show overlap of CENP-A and CENP-B following auxin treatment. Each image overlaps the signals of the CENP-A (green) and CENP-B (magenta) signals of the images to the left, at the centromere, at 3x its original sizing to demonstrate the overlap or comparative location of CENP-A and CENP-B. Unstretched chromosomes shown in the top panels and stretched chromosomes shown in bottom panels similar to the numbering scheme in (A). Scale bar 2 μm long, bottom, in grey.

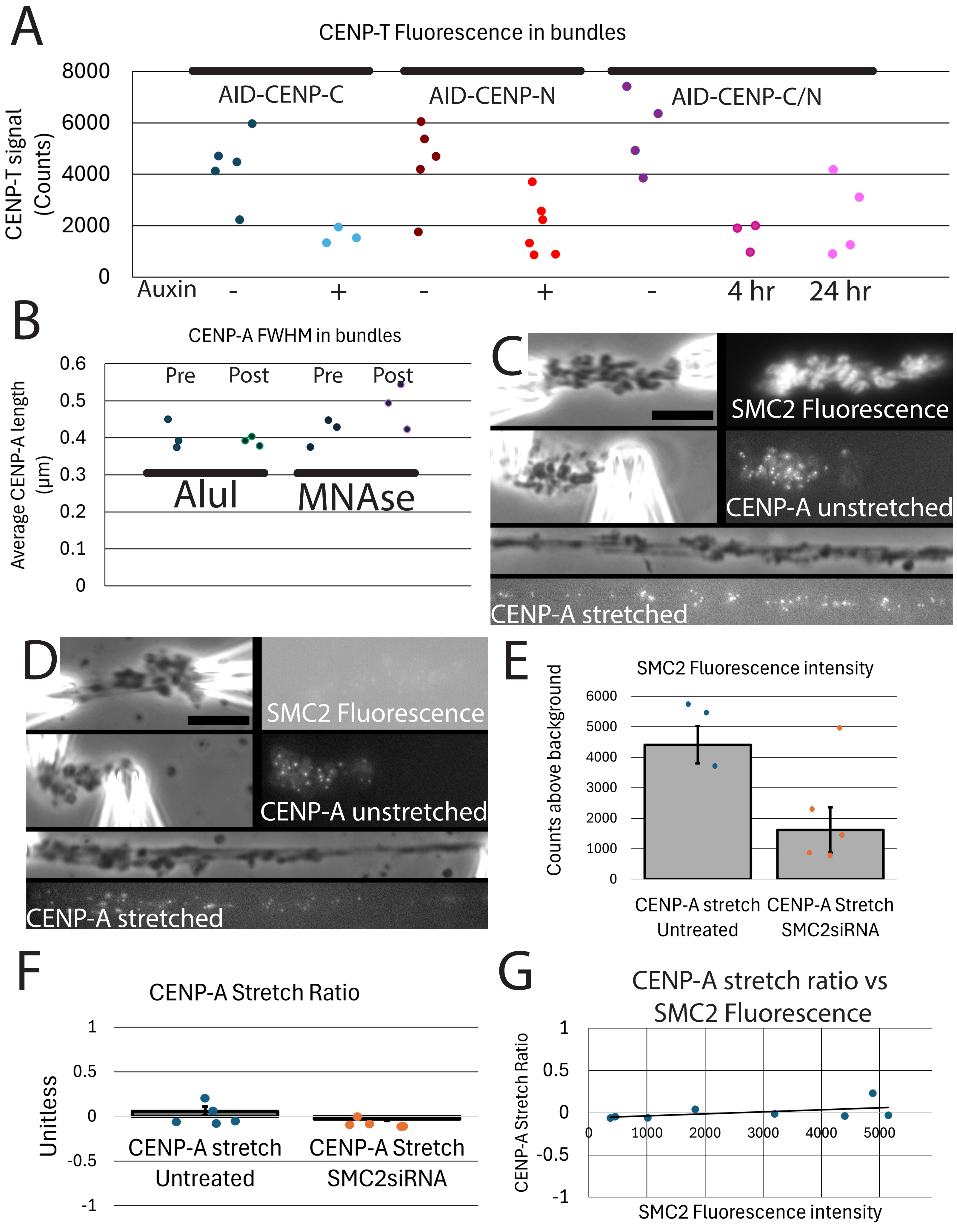
A
CENP-T Fluorescence in bundles
CENP-T signal (Counts)
8000
6000
4000
2000
0
AID-CENP-C
AID-CENP-N
AID-CENP-C/N
Auxin
-
+
-
+
-
4 hr
24 hr
B
CENP-A FWHM in bundles
Average CENP-A length (µm)
0.6
0.5
0.4
0.3
0.2
0.1
0
Pre
Post
Pre
Post
AluI
MNAse
C
SMC2 Fluorescence
CENP-A unstretched
CENP-A stretched
D
SMC2 Fluorescence
CENP-A unstretched
CENP-A stretched
E
SMC2 Fluorescence intensity
Counts above background
6000
5000
4000
3000
2000
1000
0
CENP-A stretch Untreated
CENP-A Stretch SMC2siRNA
F
CENP-A Stretch Ratio
Unitless
1
0.5
0
-0.5
-1
CENP-A stretch Untreated
CENP-A Stretch SMC2siRNA
G
CENP-A stretch ratio vs SMC2 Fluorescence
CENP-A Stretch Ratio
1
0.5
0
-0.5
-1
0
1000
2000
3000
4000
5000
SMC2 Fluorescence intensity

Supplementary Figure 6. CCAN perturbations, Nuclease, and SMC2 siRNA centromere stretching.

(A) Single Data Points for CENP-T bundle fluorescence. The average graphs from Figure 6B as single data points instead of average with error bars. Each data point and N in Figure 6B represents an average of 5 data points from a single bundle. In short, all auxin treatments caused a statistically significant decrease in CENP-T fluorescence.

(B) Single Data Points for CENP-A initial length. The average graphs from Figure 6D as the single data points instead of average with error bars. Each data point and N in Figure 6D represents an average of 5 data points from a single bundle. In short, neither AluI digestion, nor MNase digestion caused a statistically significant difference in CENP-A length.

(C) Example untreated chromosome bundles after SMC2 fluorescence staining with CENP-A stretching. Top- Chromosome bundle after spraying with fluorescent antibodies against SMC2. Top Left- bundle in phase contrast. Top Right- bundle in SMC2 fluorescence channel. Middle- Centromere lengths at initial bundle length. Middle Left- bundle in phase contrast. Middle Right- Bundle in CENP-A fluorescence channel. Bottom- Bundle after stretching. Top stretched image- Phase contrast of the stretched bundle. Bottom stretched image- CENP-A fluorescence channel. Scale bar 10 μm long, top middle, in black.

(D) Example SMC2siRNA-treated chromosome bundles after SMC2 fluorescence staining with CENP-A stretching. Top- Chromosome bundle after spraying with fluorescent antibodies against SMC2. Top Left- bundle in phase contrast. Top Right- bundle in SMC2 fluorescence channel. Middle- Centromere lengths at initial bundle length. Middle Left- bundle in phase contrast. Middle Right- Bundle in CENP-A fluorescence channel. Bottom- Bundle after stretching. Top stretched image- Phase contrast of the stretched bundle. Bottom stretched image- CENP-A fluorescence channel. Scale bar 10 µm long, top middle, in black.

(E) SMC2 fluorescence intensity of bundles with and without SMC2siRNA treatment. We traced over the chromosome bundle in phase contrast, then averaged the fluorescence intensity in the SMC2 channel with 500ms exposure time after spraying with fluorescent antibodies against SMC2. We then subtracted the general fluorescence background. The untreated bundles showed an average intensity of 4110+/-610 (N=3) counts above background, while bundles following SMC2siRNA showed an average intensity of 1620+/-720 (N=5) counts above background (statistically significantly lower compared to untreated bundles). Some bundles were not imaged at 500ms or were destroyed following fluorescent spraying, explaining the discrepancy in numbers between fluorescent measurements and stretch ratio for untreated bundles.

(F) Ratio of centromere stretching with and without SMC2 siRNA treatment. The stretch ratio of each bundle was taken by dividing the average of 5 centromeres in the stretched bundle by the average of 5 centromeres in the unstretched bundles. This was repeated 5 times for untreated bundles and repeated for 5 bundles following SMC2siRNA treatment for at least 24 hours. The untreated bundle stretch ratio was 0.056+/-0.049 (N=5) and -0.033+/-0.019 (N=5) (statistically insignificantly different) in SMC2siRNA-treated bundles.

(G) Data points and slope of all CENP-A stretch ratios plotted against SMC2 intensity. We combined the data from the untreated and SMC2siRNA data and plotted the intensity of each (E) against the CNEP-A stretch ratio (F). The slope was $2.4*10^{-5}$ stretch ratio per fluorescent count above background with an $R^2$ value of 0.240, leading us to conclude that there was very little to no correlation between SMC2 fluorescence intensity and CENP-A stretch ratio.

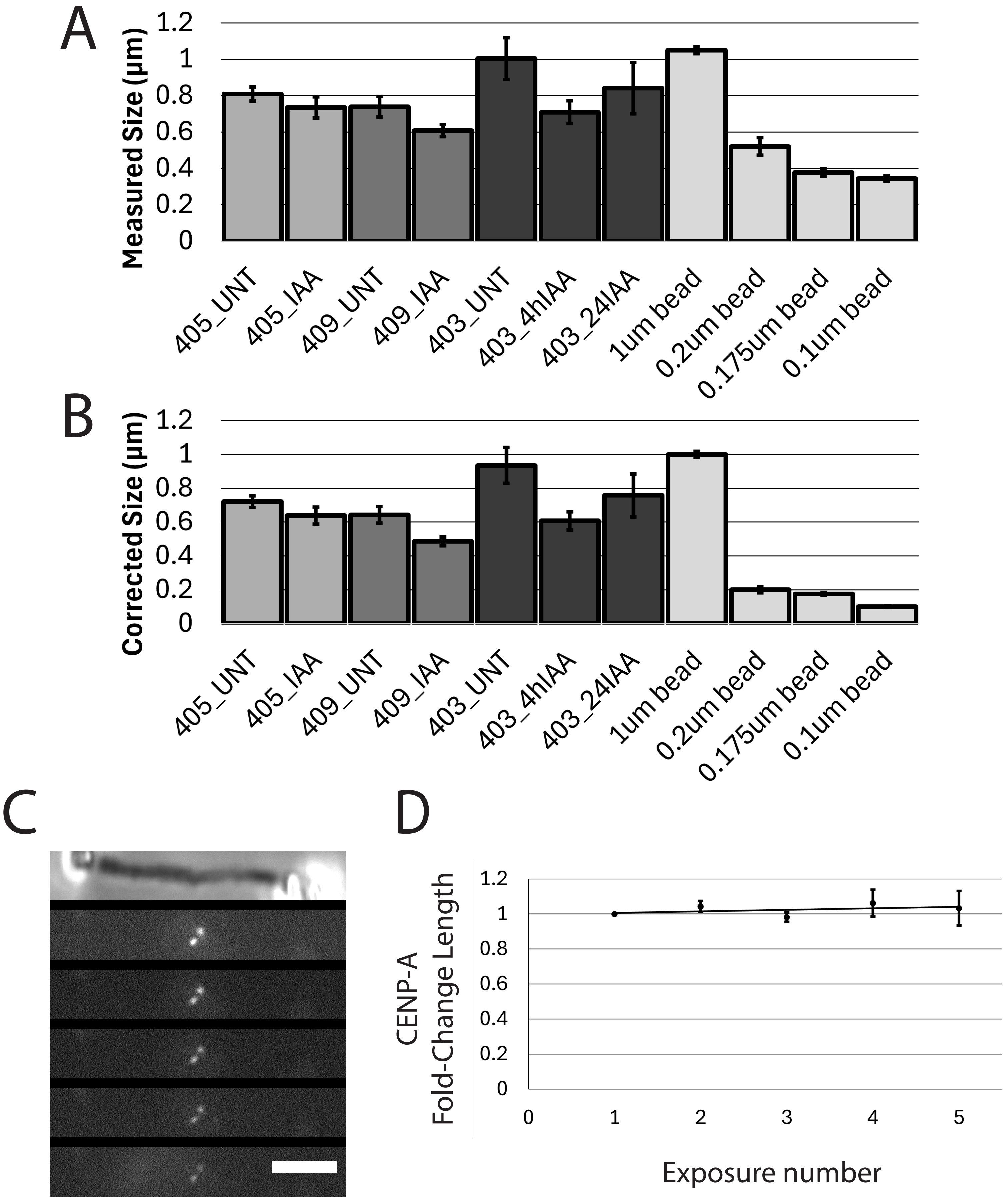
A
Measured Size (µm)
1.2
1
0.8
0.6
0.4
0.2
0
405_UNT
405_IAA
409_UNT
409_IAA
403_UNT
403_4hIAA
403_24IAA
1um bead
0.2um bead
0.175um bead
0.1um bead
B
Corrected Size (µm)
1.2
1
0.8
0.6
0.4
0.2
0
405_UNT
405_IAA
409_UNT
409_IAA
403_UNT
403_4hIAA
403_24IAA
1um bead
0.2um bead
0.175um bead
0.1um bead
C
D
CENP-A
Fold-Change Length
1.2
1
0.8
0.6
0.4
0.2
0
0
1
2
3
4
5
Exposure number

Supplementary Figure 7. Point-spread function correction and punctate bleaching tests

(A) Measured size of CENP-A and known-size beads. We imaged and processed CENP-A and known-size beads (see Materials and Methods for the specific products) through the full-width at half-maximum script to obtain their lengths when in their fluorescent channels. The measured length of CENP-A was 0.808 +/- 0.039 µm in the untreated CENP-C cell line (405_UNT), 0.735 +/- 0.058 µm in the 4-hour auxin-treated CENP-C cell line (405_IAA), 0.739 +/- 0.056 µm in the untreated CENP-N cell line (409_UNT), 0.608 +/- 0.033 µm in the 4-hour auxin-trated CENP-N cell line (409_IAA), 1.004 +/- 0.114 µm in the untreated CENP-C/N cell line (403_UNT), 0.841 +/- 0.063 µm in the 4-hour auxin-treated CENP-C/N cell line (403_4hIAA), and 0.841 +/- 0.141 µm in the 24-hour auxin-treated CENP-C/N cell line (403_24hIAA). The measured size of the known-size beads were 1.050 +/- 0.019 µm for the 1.0 µm size beads (N=15), 0.520+/- 0.048 µm for the 0.2 µm size beads (N=15), 0.377 +/- 0.020 µm for the 0.175 µm size beads (N=20), and 0.344 +/- 0.013 µm for the 0.1 µm size beads (N=15), where N are the number of beads counted.

(B) Corrected size of CENP-A and known-size beads. Knowing the size of the beads and the measured size of the beads with the microscope, we can derive the point-spread function of the microscope such that $M^2 = (P^2+A^2)$, where M is the measured size of the particle, P is the point spread function of the microscope, and A is the actual/true size of the particle, thus $P = (M^2-A^2)^{0.5}$. Using the formula, the point spread function of the microscope is 0.319+/-0.006 µm from the 1.0 µm bead, 0.480+/-0.045 µm from the 0.2 µm bead, 0.334+/-0.012 µm from the 0.175 µm beads, and 0.329 +/- 0.019 µm from the 0.1 µm beads. This gave an average point-spread function of 0.365 µm, from which all measured lengths were corrected and reported as $A=(M^2-0.365^2)^{0.5}$ µm. The corrected values for CENP-A and CENP-B can be found in Figures 4B, 4D, and Supplementary Figure 3B.

(C) Example image of CENP-A signal after repeated exposures. A chromosome was isolated and repeatedly exposed to the length of the light intensity and duration used in imaging CENP-A. The images were then analyzed using the full width at half-maximum script to obtain the initial length and following lengths of the centromere to determine the effect of measured length from repeated exposures. The upper centromere had an average slope of -0.012 change in length from its initial length per exposure, while the bottom centromere had an average slope of -0.041 change in length per exposure.

(D) Averaged graph of measured length over initial length per exposure. Each centromere in the data set (12 centromeres, from 10 chromosomes) was imaged 5 times and the length of the measured centromere was divided by its initial length. This value was then lined up with its exposure number and a slope was taken and averaged along all centromeres. The resulting slope showed an increase of length 0.008 relative to its initial length per image captured with an $R^2$ value of 0.165. Due to the small increase in relative length of the centromere per exposure and the low $R^2$ value/inconsistency in the slope, we did not make corrections to any lengths in relation to exposure number.
